# Expression of ALS-PFN1 impairs vesicular degradation in iPSC-derived microglia

**DOI:** 10.1101/2023.06.01.541136

**Authors:** Salome Funes, Del Hayden Gadd, Michelle Mosqueda, Jianjun Zhong, Jonathan Jung, Shankaracharya, Matthew Unger, Debra Cameron, Pepper Dawes, Pamela J. Keagle, Justin A. McDonough, Sivakumar Boopathy, Miguel Sena-Esteves, Cathleen Lutz, William C. Skarnes, Elaine T. Lim, Dorothy P. Schafer, Francesca Massi, John E. Landers, Daryl A. Bosco

## Abstract

Microglia play a pivotal role in neurodegenerative disease pathogenesis, but the mechanisms underlying microglia dysfunction and toxicity remain to be fully elucidated. To investigate the effect of neurodegenerative disease-linked genes on the intrinsic properties of microglia, we studied microglia-like cells derived from human induced pluripotent stem cells (iPSCs), termed iMGs, harboring mutations in profilin-1 (PFN1) that are causative for amyotrophic lateral sclerosis (ALS). ALS-PFN1 iMGs exhibited lipid dysmetabolism and deficits in phagocytosis, a critical microglia function. Our cumulative data implicate an effect of ALS-linked PFN1 on the autophagy pathway, including enhanced binding of mutant PFN1 to the autophagy signaling molecule PI3P, as an underlying cause of defective phagocytosis in ALS-PFN1 iMGs. Indeed, phagocytic processing was restored in ALS-PFN1 iMGs with Rapamycin, an inducer of autophagic flux. These outcomes demonstrate the utility of iMGs for neurodegenerative disease research and highlight microglia vesicular degradation pathways as potential therapeutic targets for these disorders.

## INTRODUCTION

Microglia are resident macrophages and primary phagocytes of the central nervous system (CNS). The phagocytic properties are critical for microglia function, including synapse pruning, circuit remodeling and clearing debris^1^. While microglia serve to sculpt and protect the CNS microenvironment, microglia dysregulation is pathologically associated with multiple neurodegenerative diseases, including Alzheimer’s disease, Parkinson’s disease, amyotrophic lateral sclerosis (ALS) and frontotemporal dementia (FTD)^2^. Transcriptomic analyses indicate that microglia adopt an altered state in the context of neurodegeneration^3^. Further, the functional properties of microglia change in a neurodegenerative disease context, wherein microglia can exhibit increased production of inflammatory mediators and altered phagocytic behaviors that are, in turn, neurotoxic^2^. However, the extent to which microglial dysfunction occurs as a response to a diseased CNS environment versus from cell-autonomous factors is not fully understood.

Rodent models have been integral for elucidating the physiological and pathological functions of microglia. However, inter-species variations often limit the biological relevance of microglia-specific findings for human disease^1^. Several protocols have recently emerged for the differentiation of human induced pluripotent stem cells (iPSCs) into microglia-like cells^4–6^, which represent a valuable cellular model to study microglia processes related to human development and neurodegenerative disease. Human microglia-like cells also represent an ideal system for investigating whether disease-linked genes modulate the intrinsic properties of microglia. For the current study, we used our optimized protocol that mimics microglia Myb-independent ontogeny for the generation of human iPSC-derived microglia-like cells, which we refer to as iMGs, in sufficient yield for downstream gene-expression and functional analyses^7, 8^. Notably, iMGs produced from our protocol express microglia-enriched markers, exhibit a transcriptome that resembles primary human microglia, respond to immune stimulation, and exhibit phagocytosis.

Herein, we used iMGs to investigate the role of profilin 1 (PFN1) in neurodegeneration. PFN1 plays an essential role in modulating actin dynamics through interactions with actin, proteins enriched with poly-L-proline motifs and phosphoinositide (PIP) lipids^9^. Autosomal dominate mutations in PFN1 are causative for the fatal neurological disorder ALS^10^, although the mechanism underlying PFN1-medidated ALS has not been elucidated. While ALS is primarily classified by upper and lower motor neuron degeneration, microglial pathology is commonly observed in ALS patients and animal models^11^. Intriguingly, *PFN1* transcripts are more abundant in microglia than neurons^12^, raising the possibility that PFN1 plays important physiological roles in microglia. In support of this notion, PFN1 becomes upregulated in reactive microglia following brain injury^13^ and PFN1 knockdown attenuates the microglia pro-inflammatory state in models of ischemia^14^. PFN1 was also identified as a modifier of phagocytosis through a CRISPR-screen in human microglia-like cells^15^. In light of these observations, we sought to determine whether ALS-linked mutations in PFN1 alter the intrinsic properties of iMGs.

Our proteomic and transcriptomic analyses of ALS-PFN1 iMGs revealed differentially expressed proteins and genes, respectively, related to lipid metabolism and vesicular degradation pathways compared to control iMGs. Vesicular degradation pathways coordinate packaging of intracellular and extracellular substrates into vesicles, which ultimately fuse with the lysosome for degradation of cargo^16^. Although ALS-PFN1 iMGs were able to engulf synaptosomes and other substrates, mutant iMGs were deficient in processing phagocytosed material through the endo-lysosomal pathway. Deficits in the autophagy pathway may underlie inefficient phagocytosis in ALS-PFN1 iMGs, which exhibited reduced conversion of microtubule-associated protein light chain 3 (LC3) from its soluble form (LC3I) to its lipidated form (LC3II). ALS-PFN1 iMGs also exhibited enhanced expression of the autophagy receptor sequestosome 1 (SQSTM1/p62) and TBC1D15, a member of the Tre2/Bub2/Cdc16 (TBC) domain family that is involved in selective autophagy^17^. Nuclear magnetic resonance (NMR) experiments revealed that ALS-PFN1 exhibits enhanced binding affinity for PI3P, a phospholipid that plays a critical role in phagosome maturation and autophagy^18, 19^, thus implicating a gain of function for mutant PFN1 with respect to phospholipid signaling in autophagy. Notably, induction of autophagic flux with Rapamycin restored phagocytic processing in ALS-PFN1 iMGs to that of control iMGs. Therefore, targeting the endo-lysosomal and autophagy pathways within microglia, in addition to neurons, may be viable therapeutic strategies for ALS and related neurodegenerative disorders.

## RESULTS

### Generation and characterization of human ALS-PFN1 iPSC-derived microglia-like cells (iMGs)

ALS-linked C71G^+/-^ and M114T^+/-^ mutations were introduced into the *PFN1* locus of the human iPSC KOLF2.1J line using CRISPR/Cas9 gene-editing^20, 21^. In addition, the homozygous M114T^+/+^ variant was created as an experimental line to investigate mutant-gene dosage on potential phenotypes (**Sup Fig. 1**). Mutant iPSCs and their respective WT isogenic control lines were differentiated into iMGs using previously published protocols with some modifications (**Figure 1a**)^6, 8^. Briefly, iPSCs were differentiated into embryoid bodies (EBs). After 21-28 days, EBs produced primitive macrophage precursors (PMPs) that were terminally differentiated into iMGs for 10 to 12 days^5, 6^. As for other reports of iPSC-derived microglia, iMGs exhibited a microglia-like morphology with some ramifications and a relatively small cytoplasmic area^4, 5^. Immunofluorescence analyses determined that >90% of iMGs expressed the microglia signature proteins purinergic receptor P2RY12 and transmembrane protein 119 (TMEM119), as well as the myeloid marker protein ionized calcium-binding adapter molecule 1 (IBA1) (**Figure 1b** **and Sup Fig. 2a**). Expression of the “homeostatic” microglia-signature genes G protein-coupled receptor 34 (*GPR34*), protein S (*PROS1*), *P2RY12* and MER proto-oncogene tyrosine kinase (*MERTK*)^22^ significantly increased in iMGs compared to PMPs and iPSCs as determined by qPCR. Additionally, expression of the myeloid transcription factor PU.1 gene (*SPI-1*) was higher in PMPs relative to iMGs, but negligible in iPSCs. Expression of the pluripotency marker SRY-Box transcription factor 2 (*SOX2*) was significantly reduced in iMGs. None of the aforementioned genes were expressed differentially between WT and ALS-PFN1 iMGs (**Figure 1c** **and Table S1**).

RNA-sequencing (RNAseq) analysis was also performed on mutant PFN1 and WT iMGs derived from five different iPSC lines, including PFN1 C71G^+/-^, M114T^+/-^, M114T^+/+^ and two WT lines (**Sup. Figure 1a**) and compared to publicly available RNASeq data sets from other studies of iPSC-derived microglia^4, 5^. Principal component analysis indicated that the transcriptome of our iMGs most closely resemble primary human microglia and were most divergent from iPSCs and monocytes (**Figure 1d**). We also evaluated genes that reportedly change as a function of microglia developmental stage^23^ (**Sup Fig. 2b**). Read counts for genes associated with adult microglia, including V-Maf musculoaponeurotic fibrosarcoma oncogene homolog B (*MAFB*), monocyte differentiation antigen CD14 (*CD14*), and the colony-stimulating factor-1 receptor *(CSFR1)* were higher than for genes associated with fetal microglia, such as minichromosome maintenance complex component 5 (*MCM5*). Only one gene associated with yolk-sac microglia progenitors (coagulation factor XIII A chain, *F13A1*) was detected in iMGs and, as expected, with low abundance^23^. Notably, abundant levels of *APOE* and *SPP1*, genes that are highly expressed in aged microglia, were detected in both mutant PFN1 and WT iMGs^24^.

The reactivity of iMGs to lipopolysaccharide (LPS) was then assessed by quantifying cytokine release. At baseline, WT and PFN1 C71G^+/-^ iMGs secreted low levels of the pro-inflammatory cytokines interleukin (IL)-6, tumor necrosis factor-alpha (TNF-α) and regulated on activation normal T cell expressed and secreted (RANTES or CCL5) as well as the anti-inflammatory cytokine IL-10. The levels of IL6, CCL5, and IL10 increased significantly 24h-post LPS treatment, whereas TNF-α levels were significantly elevated after 6h of LPS stimulation (**Figure 1e** **and Table S1**). The cytokine levels were similar between mutant and WT PFN1 cells and confirmed that iMGs from both genotypes are responsive to immune stimulation.

**Figure 1.**
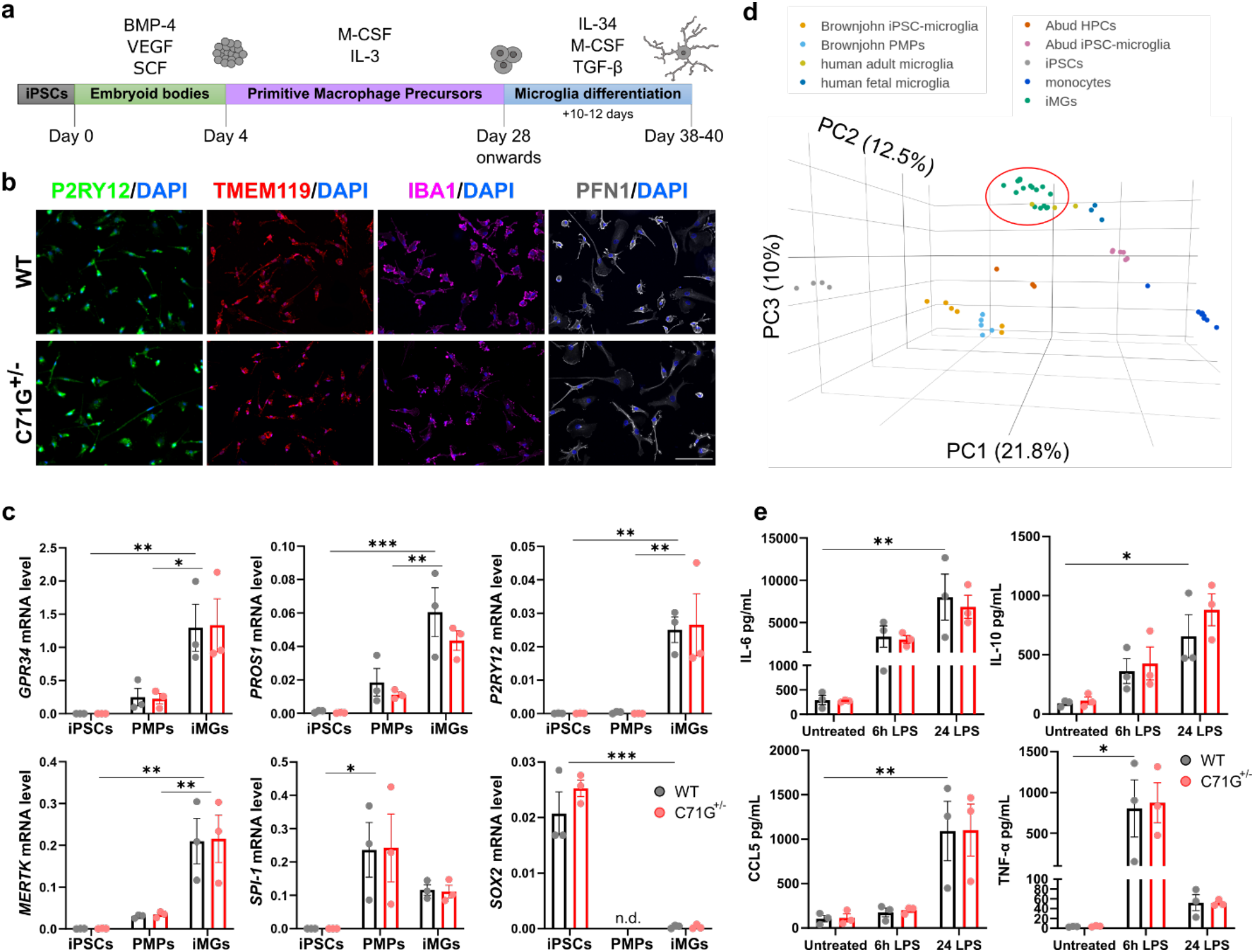
Generation and characterization of WT and ALS-PFN1 iPSC-derived microglia-like cells (iMGs). **a**, Schematic representation of iMG differentiation protocol. Bone morphogenetic protein 4 (BMP-4), vascular endothelial growth factor (VEGF), and stem cell factor (SCF) are used for embryoid body differentiation. Macrophage colony-stimulating factor (M-CSF) and interleukin (IL)-3 are included during PMP generation. IL-34, M-CSF and transforming growth factor beta (TGF-β) are added during terminal iMG differentiation. **b,c,** WT and C71G^+/-^ iPSCs differentiated into iMGs. **b**, Representative immunofluorescence images of P2RY12, TMEM119, and PFN1. Scale bar: 100 µm. **c,** Comparison of gene expression levels between iPSCs, PMPs and iMGs of myeloid and microglia-enriched genes including *GPR34* (\**P*=0.0173, ** *P*=0.0044), *PROS1* (\*\**P*=0.0040, \*\*\**P*=0.0002), *P2RY12* (\*\**P*=0.0030 for iPSC vs iMG, \*\**P*=0.0033 for PMP vs iMG), *MERTK* (\*\**P*=0.0016 for iPSC vs iMG, \*\**P*=0.0049 for PMP vs iMG), and *SPI-1* (\**P*=0.0240) and the pluripotency marker *SOX2* (\*\*\**P*=0.0003) **d,** Three dimensional principal component analysis (PCA) of iMGs from this study (green; WT n=7, C71G^+/-^ n=4, M114T^+/-^ n=3, M114T^+/+^ n=3, where n refers to an independent differentiation), primary human adult (yellow) and fetal microglia (navy blue) from Abud et al. 2017, iPSC-derived microglia-like cells (pink) and the intermediate hematopoietic progenitors cells (HPCs, dark orange) from Abud et al. 2017, iPSC-derived microglia-like cells (light orange) and the intermediate progenitors PMPs (light blue) from Brownjohn et al. 2018, and monocytes (dark blue) and iPSCs (grey) from Abud et al. 2017. Variance of each principal component (PC) is indicated in parenthesis along the axes. **e**, WT and C71G^+/-^ iMGs secrete elevated levels of IL-6 (\*\**P*=0.0050), IL-10 (\**P*=0.0156), CCL5 (\*\**P*=0.0077) and TNF-α (\**P*=0.0208) after 6h or 24h of 100 ng/mL LPS stimulation compared to untreated cells. Statistics were determined by two-way ANOVA and Šídák’s multiple comparisons test. No WT vs C71G^+/-^ comparisons were statistically significant. *P* values are listed only for WT cells for simplicity; all other statistical comparisons are defined in Table S1. Mean ± SEM for n=3 independent differentiations is shown for all bar graphs, with each data point representing an individual differentiation.

### Factors involved in lipid metabolism and phagocytosis are differentially expressed in ALS-PFN1 iMGs compared to controls

Next, we investigated proteins and genes that were differentially expressed between ALS-PFN1 and WT iMGs. Quantitative proteomics with 6-plex tandem mass tag (TMT) labeling of whole cell lysates was used to assess differential protein expression between PFN1 C71G^+/-^ and WT iMGs by mass spectrometry. A total of 1813 proteins in 1660 clusters were detected in the TMT analysis, with 56 proteins differentially expressed between PFN1 C71G^+/-^ and WT iMGs (**Figure 2a** **and Table S2**). Of those 56 proteins, 30 proteins were upregulated in PFN1 C71G^+/-^ iMGs including fatty acid-binding protein (FABP) 4 and FABP5. PFN1 was the most significantly downregulated protein. Consistent with previous findings in PFN1 C71G^+/-^ and M114T^+/-^ patient-derived lymphoblast cells^25, 26^, all mutant PFN1 iMGs expressed lower PFN1 levels compared to control cells by Western blot analysis, with ∼60% reduction in PFN1 C71G^+/-^ iMGs (**Figure 2b,c**) and ∼30% reduction in PFN1 M114T^+/-^ iMGs (**Figure 2b,d**). Strikingly, PFN1 levels were reduced by 80% in M114T^+/+^ iMGs compared to WT iMGs (**Figure 2b,d**). These results are consistent with a destabilizing effect of ALS-linked mutations on PFN1 structure, leading to enhanced proteasomal degradation at steady-state^25^.

Metascape was used for functional enrichment analysis of the 56 differentially expressed proteins. The KEGG pathway “ferroptosis”, a form of iron-dependent cell death caused by accumulation of lipid peroxides^27^, was the most significantly enriched term and the “fatty acid transporter” pathway was among the top 20 (**Figure 2e** **and Table S3**). Additional lipid-related terms, including the cellular component “lipid droplet” (**Sup. Fig 3a and Table S4**) and the molecular function “cholesterol binding” (**Sup. Fig 3b and Table S4**), were among the top enriched gene ontology (GO) terms from an Enrichr analysis. In light of these lipid-related terms, we probed for evidence of lipid accumulation in the form of lipid droplets. Lipid droplets contain a core of neutral lipids surrounded by a phospholipid monolayer. While lipid droplets represent lipid storage sites under physiological conditions, aberrant accumulation of lipid droplets correlates with aging and disease^28, 29^. Mutant PFN1 and WT iMGs were stained with BODIPY (**Figure 2f,g**), a fluorescent dye that binds lipid droplets, and the BODIPY signal was quantified by fluorescence microscopy. Mutant PFN1 iMGs presented with more BODIPY signals compared to WT iMGs, especially in PFN1 C71G^+/-^ (**Figure 2h**) and M114T^+/+^ (**Figure 2i**) iMGs. A low percentage (<5%) of both ALS-PFN1 and WT iMGs exhibited a dense area of lipid droplets, which appeared to fill most of the cell. Upon co-staining for BODIPY and anti-PFN1, PFN1 signal was clearly observed surrounding lipid droplets in these cells (**Figure 2j**). While PFN1 may also be associated with lipid droplets in cells presenting with fewer lipid droplets, we note that detection of PFN1 surrounding lipids droplets was most obvious in iMGs with high lipid droplet content. These observations raise the possibility that PFN1 associates with lipid droplets, potentially through direct binding of PFN1 with the phospholipid outer shell.

**Figure 2.**
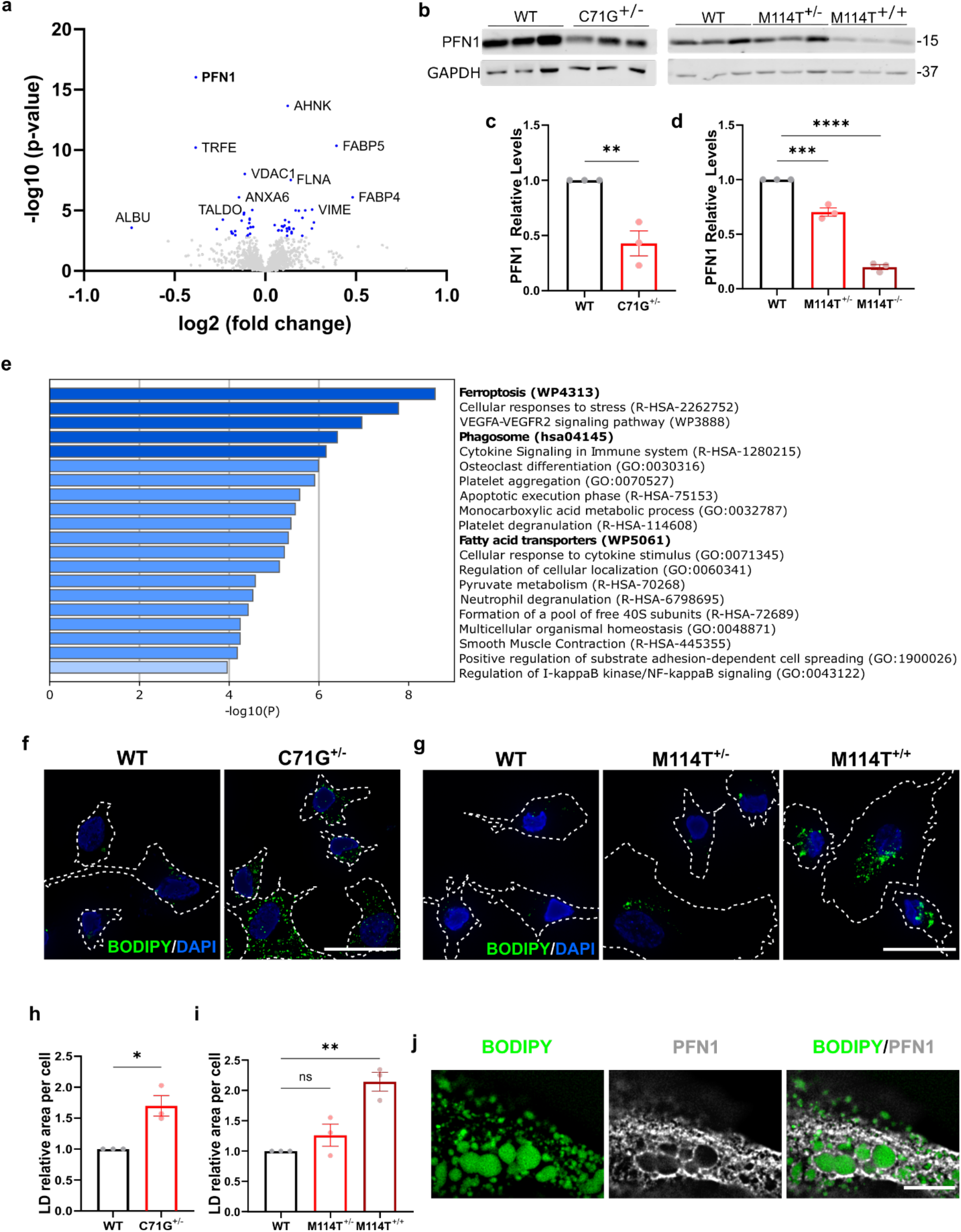
Quantitative proteomics analysis highlights dysregulation of lipid metabolism and phagocytosis in ALS-PFN1 iMGs. All data include n=3 independent differentiations. Bar graphs show mean ± SEM with individual data points representing independent differentiations. **a,** Volcano plot of proteins differentially expressed between C71G^+/-^ and WT iMGs identified by TMT proteomics. Log2 fold change in expression and significance (-log10 p-value) are displayed on the x- and y-axes, respectively. Differentially expressed proteins with a *P* value <0.00160 (blue dots) are considered significant after the Benjamini-Hochberg test (see **Supp. Table 2**). **b,** Western blot analysis of PFN1 levels in C71G^+/-^, M114T^+/-^, M114T^+/+^ iMGs compared to control WT iMGs. **c,d,** Quantification of **b**. For each independent differentiation, PFN1 protein levels were normalized first to GAPDH (loading control) and then to the respective WT control from the same differentiation. Statistics were determined using unpaired two-tailed t-test for **c** (\*\**P*=0.0073, t=5.035, df=4) and ordinary one-way ANOVA with Dunnett’s multiple comparisons test for **d** (\*\*\**P*=0.0003, q=8.232, DF=6, and \*\*\*\**P*<0.0001, q=22.20, DF=6). **e,** Heatmap of enriched terms across differentially expressed proteins generated by Metascape (see **Supp. Table 3**). The bars are colored-coded according to *P* value, with the significance (-log *P*) of the term defined on the x-axis. Terms of interest are emboldened. **f-j**, Lipid droplets accumulate in mutant PFN1 iMGs as determined by BODIPY immunofluorescence analysis. **f,g,** Representative immunofluorescence images of BODIPY staining in **(f)** PFN1 WT and C71G^+/-^ iMGs and **(g)** PFN1 WT, M114T^+/-^ and M114T^+/+^ iMGs. Cell boundaries are depicted with white dashed lines. Scale bar= 25µm. **h,i,** Quantification of the area of BODIPY fluorescence signal representing lipid droplets (LD) was normalized to WT iMGs within each independent differentiation for **f** (unpaired two-tailed t-test, \**P*=0.0137, t=4.201, df=4) and **g** (ordinary one-way ANOVA and Dunnett’s multiple comparisons test, ns *P*=0.3647, q= 1.338, DF=6 for WT vs M114T^+/-^ and \*\**P*=0.0020, q=5.840, DF=6 for WT vs M114T^+/+^). **j,** Representative immunofluorescence images (one Z-plane acquired by focusing on prominent lipid droplets) showing PFN1 (gray) surrounding lipid droplets (BODIPY in green) in C71G^+/-^ iMGs. Scale bar=10µm.

A differential gene expression analysis was also performed with RNASeq data from “ALS-PFN1” iMGs, including both PFN1 C71G^+/-^ and M114T^+/-^ lines, compared to the isogenic WT lines. Although only thirteen genes were differentially expressed with statistical significance, thus precluding a pathway analysis, *TBC1D15* emerged as a highly significant, upregulated gene in ALS-PFN1 iMGs (**Figure 3a** **and Table S5**). TBC1D15 is a GTPase-activating protein for RAB7 and is involved in autophagosome biogenesis during selective autophagy^17^. Elevated *TBC1D15* mRNA in ALS-PFN1 iMGs was validated by qPCR (**Figure 3b,c**). Further, TBC1D15 upregulation at the protein level in ALS-PFN1 iMGs was confirmed through Western blot (**Figure 3d,e**) and immunofluorescence (**Figure 3f,g**) analyses. Conversely, no difference in Rab7 levels were detected between WT and mutant PFN1 iMGs (**Sup. Figure 4**). TBC1D15 staining resulted in the expected punctate pattern with a vesicular morphology in both WT and PFN1 C71G^+/-^ iMGs (**Figure 3f**) with significantly more TBC1D15 signal in mutant iMGs (**Figure 3g**)^30^. Notably, TBC1D15 and PFN1 colocalized to a greater extent in C71G^+/-^ iMGs than WT iMGs according to the Pearson’s correlation analysis (**Figure 3h**). While a similar fraction of TBC1D15 signal co-occured with PFN1 between genotypes according to the Mander’s coefficient (**Figure 3i**), a higher fraction of PFN1 signal co-occurred with the TBC1D15 signal in C71G^+/-^ iMGs (**Figure 3j**). Moreover, PFN1 signal prominently colocalized with TBC1D15 punctuated structures as demonstrated by fluorescence signal intensity plots (**Figure 3k-m**). Given that the total levels of PFN1 are lower in C71G^+/-^ iMGs compared to controls (**Figure 2b-d**), these outcomes indicate that a larger fraction of the PFN1 protein remaining in the cell colocalizes with TBC1D15 in ALS-PFN1 iMGs.

As lentiviral transduction was toxic to iMGs, we pursued PFN1 knockdown studies in the human microglia immortalized HMC3 line to assess whether TBC1D15 expression is due to loss of PFN1 function. In contrast to ALS-PFN1 iMGs, knockdown of PFN1 correlated with significantly reduced levels of TBC1D15 in (**Sup Figure 5**). This outcome implicates a physiological relationship between PFN1 and TBC1D15 in the context vesicular degradation and supports the notion that mutant PFN1 exerts a gain of aberrant function in this pathway.

**Figure 3.**
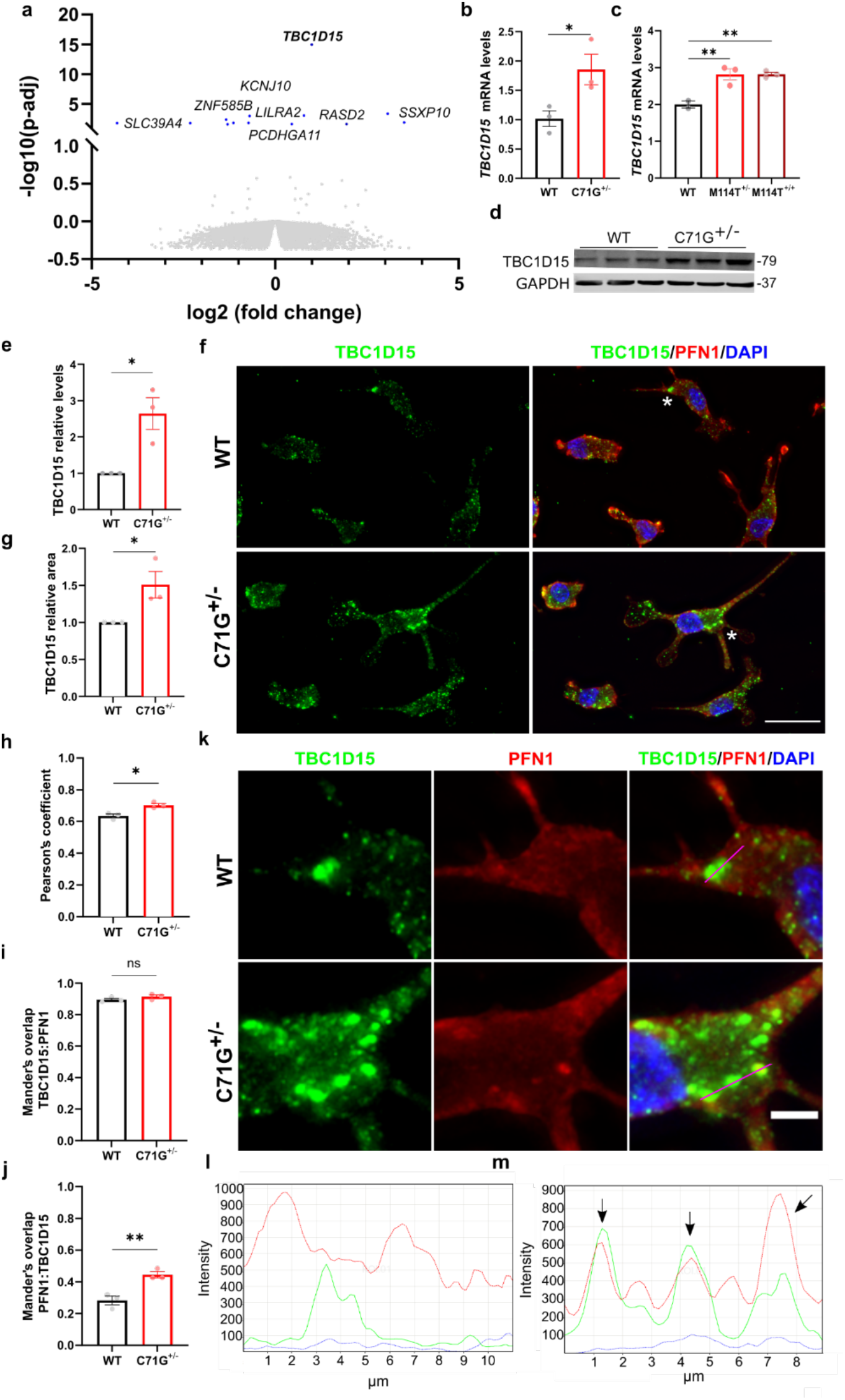
TB1C1D15 is upregulated in ALS-PFN1 iMGs and colocalizes with PFN1. **a**, Volcano plot of differentially expressed genes between ALS-PFN1 iMGs (n=4 for C71G^+/-^ iMGs and n=3 for M114T^+/-^ iMGs) relative to WT iMGs (n=7), where n refers to an independent differentiation; see **Table S5**. Differentially expressed genes (*P* adjusted value<0.05) are highlighted in blue. **b,c,** Relative mRNA levels of *TBC1D15* normalized to the average of the respective WT iMGs for n=2-3 independent differentiations (**b,** \**P*=0.0452, t=2.875, df=4; **c**, WT versus M114T^+/-^ \*\**P*= 0.0030, q=10.60, DF=3; WT versus M114T^+/+^ \*\**P*= 0.0042, q=9.403, DF=3). **d,** Enhanced TBC1D15 protein expression determined by Western blot analysis. **e,** Quantification of **d**. For each independent differentiation TBC1D15 levels were normalized to GAPDH and then to the levels of the respective WT line from the same differentiation (\**P*=0.0197, t=3.763, df=4). **f-m,** TBC1D15 colocalizes with PFN1 in WT and C71G^+/-^ iMGs **f,** Representative immunofluorescence images of TBC1D15 (green) and merged images of TBCD15, PFN1 (red) and DAPI (blue) showing regions of overlayed TBC1D15:PFN1 signal in yellow. Scale bar=50 µm. Asterisks indicate cells used for analysis in k-m. **g,** Quantification of the area of TBC1D15 signal per mutant iMG cell was normalized to WT iMGs from the same differentiation (*\*P*=0.0458, t=2.862, df=4).**h,** Pearson’s correlation coefficient of TBC1D15 and PFN1 signal (*\*P*=0.0292, t=5.722, df=2). **i,j,** Mander’s co-occurrence coefficient indicating the fraction of **(i)** TBC1D15:PFN1 co-occurrence (ns *P*=0.2214, t=1.447, df=4) and **(j)** the fraction of PFN1:TBC1D15 co-occurrence (*\*\*P*=0.0085, t=4.818, df=4). **k,** Zoomed regions of interest from **f** used to obtain fluorescent signal intensity plots. Scale bar=5 µm. **l,m,** Fluorescent signal intensity plots across the lines shown in **k** (merged) of TBC1D15 vesicle-like structures for WT **(l)** and C71G^+/-^ iMGs **(m)**. Arrows indicate overlap of TBC1D15 and PFN1 signals. All bar graphs show mean ± SEM of data derived from independent differentiations, where each data point represents an independent differentiation. Statistics were determined using unpaired two-tailed t-test for WT vs C71G^+/-^ iMGs comparisons or ordinary one-way ANOVA with Dunnett’s multiple comparisons test for WT vs M114T^+/-^ and M114T^+/+^ comparisons.

### ALS-PFN1 microglia exhibit deficits in vesicular degradation of phagocytosed material *in vitro* and *in vivo*

A critical function of microglia is phagocytosis, a process that was also highlighted by our proteomics analysis (**Figure 2e**). We examined phagocytosis in iMGs using live-cell imaging with substrates that were labeled with pHrodo red (pHrodo), a pH-sensitive dye that emits red fluorescence exclusively in low pH cellular compartments. Low pH cellular compartments include late endosomes and phagolysosomes, the latter being generated upon fusion of phagosomes and lysosomes during phagocytosis^31^. Initially, we administered pHrodo-labeled mouse synaptosomes to PFN1 C71G^+/-^ and WT iMGs, resulting in enhanced red fluorescent signals in both lines by 30 min that became more pronounced by 2h (**Figure 4a**). Quantification of a phagocytosis index for each line revealed a significant increase in pHrodo signal in PFN1 C71G^+/-^ versus WT iMGs over the course of 12h. Addition of cytochalasin D, an inhibitor of actin polymerization that prevents phagocytosis, significantly attenuated pHrodo fluorescence (**Figure 4b,c**)^32^. Similar to PFN1 C71G^+/-^, PFN1 M114T^+/-^ and M114T^+/+^ iMGs also produced enhanced pHrodo fluorescence relative to their WT counterparts in this assay. To verify that the pHrodo-labeled signal was originating from intracellular acidic compartments, we pretreated iMGs with bafilomycin A (BafA), a vacuolar H^+^-ATPase inhibitor that blocks acidification of the phagolysosome^33^. As expected, red fluorescent signal is undetected in all BafA-treated iMGs (**Figure 4d,e**). Similar phagocytosis assays were conducted with other disease-relevant substrates including human synaptosomes isolated from iPSC-derived lower motor neurons and aggregated C71G PFN1 recombinant protein. Administration of both substrates resulted in a significantly higher phagocytosis index in PFN1 C71G^+/-^ iMGs relative to WT iMGs (**Sup. Figure 6**).

Based on the phagocytosis assays with pHrodo-labeled material alone, it is unclear whether higher pHrodo signal in mutant iMGs stems from enhanced substrate uptake or inefficient substrate degradation. To distinguish between these possibilities, a synaptosome washout step was included in the phagocytosis assay, allowing for quantification of the initial uptake of synaptosomes as well as the amount of undegraded synaptic material after 48h post-washout (**Figure 4f**). Human synaptosomes were used here, as iMGs processed these more rapidly than murine synaptosomes. To track synaptosomes in the initial phase of the assay, iMGs were incubated with constitutively fluorescent synaptosomes for 15 minutes, after which unbound material was washed away (referred to as 0h post-washout). Alexa fluor-labeling was used instead of pHrodo, as pHrodo cannot be used to monitor phagocytosed material before it enters acidic compartments. An engulfment index was calculated based on the area of synaptosomes in direct contact with and inside iMGs, as delimitated by anti-IBA1 staining (**Figure 4g**) and was found to be similar for PFN1 C71G^+/-^ and WT iMGs (**Figure 4h**). Given that actin polymerization is required during the early stages of phagocytosis and that ALS-linked PFN1 variants can modify actin polymerization^25, 34^, we also measured F-actin levels with phalloidin in fixed-cells at baseline (i.e., in the absence of synaptosomes) and at 0h post-washout. A redistribution of F-actin signal was observed in both PFN1 C71G^+/-^ and WT iMGs upon addition of synaptosomes, but there were no discernable differences in F-actin signals between genotypes (**Sup. Figure 7**). Therefore, it is unlikely that deficits in synaptosome uptake or actin polymerization underlie the differences in phagocytosis between mutant PFN1 and WT iMGs.

At 48h post-washout (**Figure 4f**), the volume of pHrodo-labeled synaptosomes was quantified within compartments that stained for cluster of differentiation 68 (CD68), a macrophage/microglia-specific endo-lysosomal marker (**Figure 4i**). PFN1 C71G^+/-^ iMGs contained a significantly higher volume of synaptosomes within CD68-positive compartments compared to WT iMGs (**Figure 4j**). Further, a higher percentage of PFN1 C71G^+/-^ iMGs retained pHrodo-labeled synaptosomes compared to control iMGs (**Figure 4k**). Together, these data indicate that vesicular degradation of phagocytosed material, rather than the uptake of material, is less efficient in ALS-PFN1 iMGs compared to controls.

**Figure 4.**
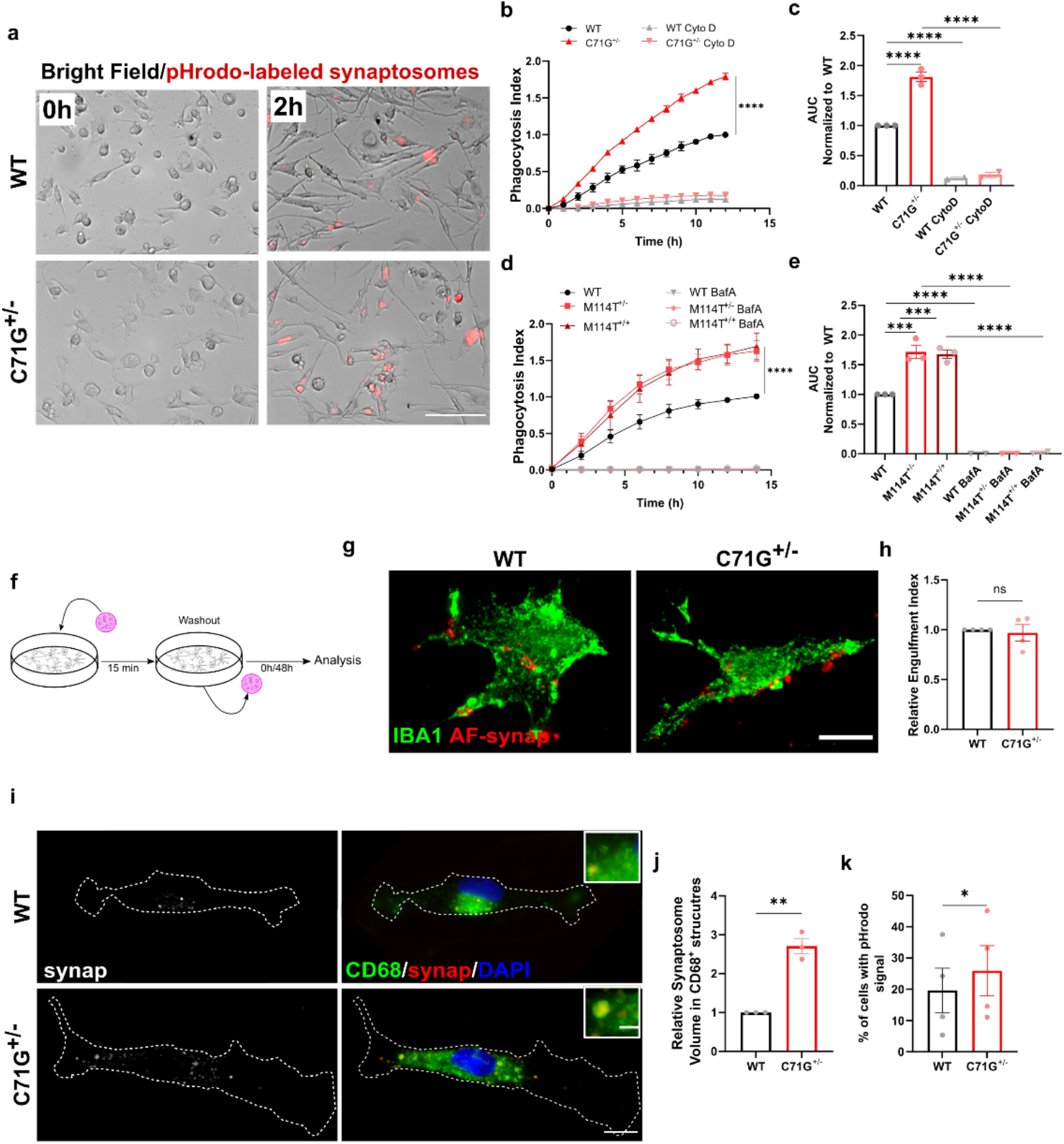
ALS-PFN1 iMGs exhibit inefficient phagocytic degradation. **a-e**, Phagocytosis of pHrodo-labeled mouse-synaptosomes by iMGs. **a,** Representative images of synaptosomes engulfed by WT and C71G^+/-^ iMGs at the indicated timepoints. Scale bar=100 µm. **b,** Quantification of the phagocytosis index (see Methods) over 12h for C71G^+/-^ compared to WT iMGs (\*\*\*\**P*<0.0001, two-way ANOVA with Tukey’s multiple comparison test, *F*(3, 78) = 1746). WT and C71G^+/-^ iMGs were pre-treated with 10 µM Cytochalasin D (CytoD). **c,** Area under the curve (AUC) determined from **b** (\*\*\*\**P*<0.0001, one-way ANOVA with Tukey’s multiple comparison test, q=17.28, DF=6 for WT vs C71G^+/-^). **d,** The same analysis as **b** with WT, M114T^+/-^ and M114T^+/+^ iMGs (****P<0.0001, two-way ANOVA with Tukey’s multiple comparison test, F (7, 72) = 36.50). iMGs were pre-treated with 100 nM bafilomycin A (BafA). **e,** AUC determined from **d** (\*\*\**P*=0.0002, q=11.29, DF=9 for WT vs M114T^+/-^ and \*\*\**P*=0.0003, q=10.71, DF=9 for WT vs M114T^+/+^). Additional statistical comparisons for **b** and **d** are in **Table S1**. **f-k,** Washout assay using human synaptosomes for WT and C71G^+/-^ iMGs. **f,** Schematic of the assay. **g,h,** Assay using synaptosomes labeled with Alexa Fluor (AF-synap) at 0h post-washout. **g,** Representative immunofluorescence images of iMGs labeled with IBA1 (green) engulfing AF-synap (red). Scale bar: 10 µm **h,** Quantification of the engulfment index (see Methods) normalized to WT iMGs (ns *P*=0.7185, unpaired two-tailed t-test, t=0.3779, df=6). **i-k,** Assay using pHrodo-labeled synaptosomes (synap) at 48h post-washout. **i,** Representative immunofluorescence images of the residual pHrodo-synaptosome signal (white in left-side images, red in right-side images) and merged with CD68 (green). Zoomed images of pHrodo-synaptosomes colocalizing with CD68 signal. Scale bar: 10 µm and 2 µm. **j,** Quantification of the fraction of CD68 volume occupied by pHrodo-synaptosome signal (yellow in **i**) normalized to WT iMGs (*\*\*P*=0.0011, unpaired two-tailed t-test, t=8.401, df=4). **k,** Quantification of the percentage of cells that retain pHrodo-synaptosome signal in each independent differentiation (*\*P*=0.0490, unpaired two-tailed t-test, t=2.462, df=6). All graphs in this figure show mean ± SEM for n=3-4 differentiations, with each data point representing an independent differentiation.

To investigate whether deficient degradation of phagocytosed material occurs in *in vivo* models of ALS-PFN1, dead neurons were stereotactically injected into the motor cortex of PFN1 C71G^+/-^ knock-in and WT mice (**Figure 5**), as microglia are known to phagocytose dead neuron material within the mammalian brain^35^. PFN1 C71G^+/-^ mice were generated with CRISPR/Cas9 technology such that the analogous C71G mutation was introduced into the endogenous mouse *PFN1* gene locus at one allele, resulting in physiological expression of PFN1 C71G. Dead neurons were co-labeled with the constitutively fluorescent dye Alexa fluor 546 (AF) to measure the total amount of material present in the brain tissue and the pH-sensitive dye Cypher5E (Cy5E) to quantify material that is specifically located within acidic phagosomal compartments (**Figure 5a,b**). Seventy-two hours post-injection, a range of AF signal was detected in the mouse brains from both genotypes; this range likely reflects some variability in the amount of dead neuron material that effectively diffused within the tissue post-injection (**Figure 5c**). Notably, a significantly higher fraction of the dead neuron material was retained in acidic compartments of PFN1 C71G^+/-^ mouse tissue compared to WT littermates (**Figure 5a,d**). Further, brain sections from PFN1 C71G^+/-^ mice that contained dead neurons exhibited higher Iba1 signals (**Figure 5e,f**). This was not due to more Iba1-postive cells, which were similar in number between genotypes except for lower numbers in the mutant mouse tissue represented by Ring 3 (**Figure 5g**). These data are consistent with heightened microglial reactivity and enhanced Iba1 expression in response to the presence of dead neurons PFN1 C71G^+/-^ mice. Collectively, these results support the notion that expression of ALS-linked PFN1 in phagocytic cells impairs degradation of engulfed substrates both *in vitro* and *in vivo*.

**Figure 5.**
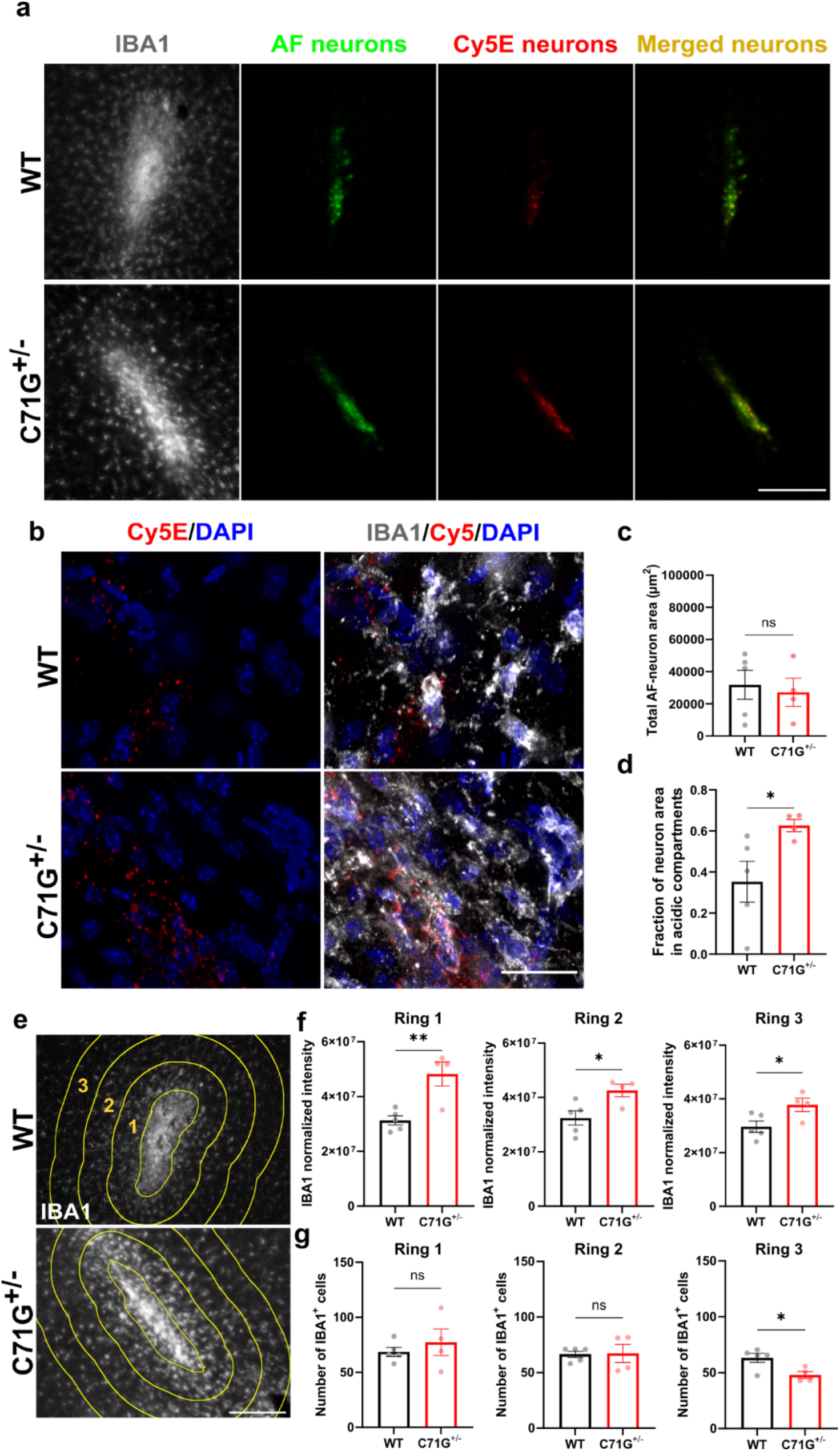
Differential processing of dead neurons in ALS-PFN1 mouse brains compared to controls. Analysis of PFN1 C71G^+/-^ (n=4) and WT (n=5) mouse brains injected with dead neurons co-labeled with the constitutively fluorescent dye Alexa fluor 546 (AF-neurons) and the pH-sensitive dye Cypher5E (Cy5E-neurons) 72h post-injection. **a,** Representative immunofluorescent images at the injection site in the mouse motor cortex of the myeloid marker IBA1(grey), the residual dead neurons (AF-neurons in green), and the dead neurons localized in acidic compartments (Cy5E-neurons in red) as well as the overlaid image of AF-neurons and Cy5E neurons (merged in yellow). Scale bar: 200 µm. **b,** Representative image of IBA1-positive (IBA1^+^)cells colocalizing with Cy5E-neuron signal from a PFN1 WT (top) and PFN1 C71G^+/-^ (bottom) mouse brain. Scale bar: 25 µm. **c,** Quantification of the total area of residual AF-neuron signal (ns *P*=0.7288, t=0.3610, df=7). **d,** The fraction of Cy5E-neuron (total area) signal that overlays with the AF-neuron (total area) signal from **c** (*\*P*=0.0499, t=2.366, df=7). **e,** Representative IBA1 (grey) immunofluorescence images at the injection site for WT PFN1 and PFN1 C71G^+/-^ mice. Four concentric rings (yellow) around the site of injection (center ring) are shown for analysis in **f** and **g**. Scale bar: 100 µm. **f,** Quantification of total IBA1 signal intensity in the three rings surrounding the injection site labeled in **e** normalized to the number of IBA^+^ cells (**P=0.0056, t=3.942, df=7 for ring 1; *P=0.0264, t=2.804, df=7 for ring 2; *P=0.0374, t=2.563, df=7 for ring 3). **g,** Quantification of the number of IBA1^+^ cells in each ring indicated in **e** (ns P=0.4752, t=0.7545, df=7 for ring 1; (ns P=0.9264, t=0.09573, df=7 for ring 2; *P=0.0255, t=2.827, df=7). Unpaired two-tailed t-test was used for all statistical comparisons. Graphs in this figure show mean ± SEM. Data points represent individual animals.

### Lysosomes accumulate in the perinuclear region of ALS-PFN1

As lysosomes represent a terminal organelle in the endo-lysosomal pathway, through which cargo is degraded during phagocytosis, endocytosis, and autophagy, we probed for evidence of lysosomal dysfunction in ALS-PFN1 iMGs. The quenched-bovine serum albumin (DQ-BSA) assay was used to measure lysosomal degradative capacity in iMGs. The DQ-BSA substrate is internalized through pinocytosis, and subsequent proteolysis by lysosomal proteases results in a fluorescence signal^36^. No significant differences in DQ-BSA signal intensity were observed between PFN1 C71G^+/-^ or M114T^+/-^ iMGs and WT controls (**Figure 6a,b**). The pH of intracellular acidic compartments within iMGs was also examined with Lysosensor DND-189. As expected, pre-treating iMGs with BafA resulted in a decrease in fluorescence, consistent with an increase in lysosomal pH (**Figure 6c**). However, there was no difference in DND-189 intensity between ALS-PFN1 iMGs and WT controls. Further, similar levels of both the lysosomal-associated membrane protein 1 (LAMP1) and the mature lysosomal hydrolase Cathepsin D (mCTSD) were detected in PFN1 C71G^+/-^ and WT iMGs by Western blot analysis (**Figure 6d-f**). Collectively, these data argue against inherent defects in lysosome function in ALS-PFN1 iMGs. Strikingly, perinuclear clustering of LAMP1-positive lysosomes was observed in approximately twice as many ALS-PFN1 iMGs than WT iMGs (**Figure 6g,h**). Excessive perinuclear clustering has been observed when autophagic flux is blocked^37^, suggesting ALS-linked PFN1 expression may perturb the autophagy pathway in iMGs.

**Figure 6.**
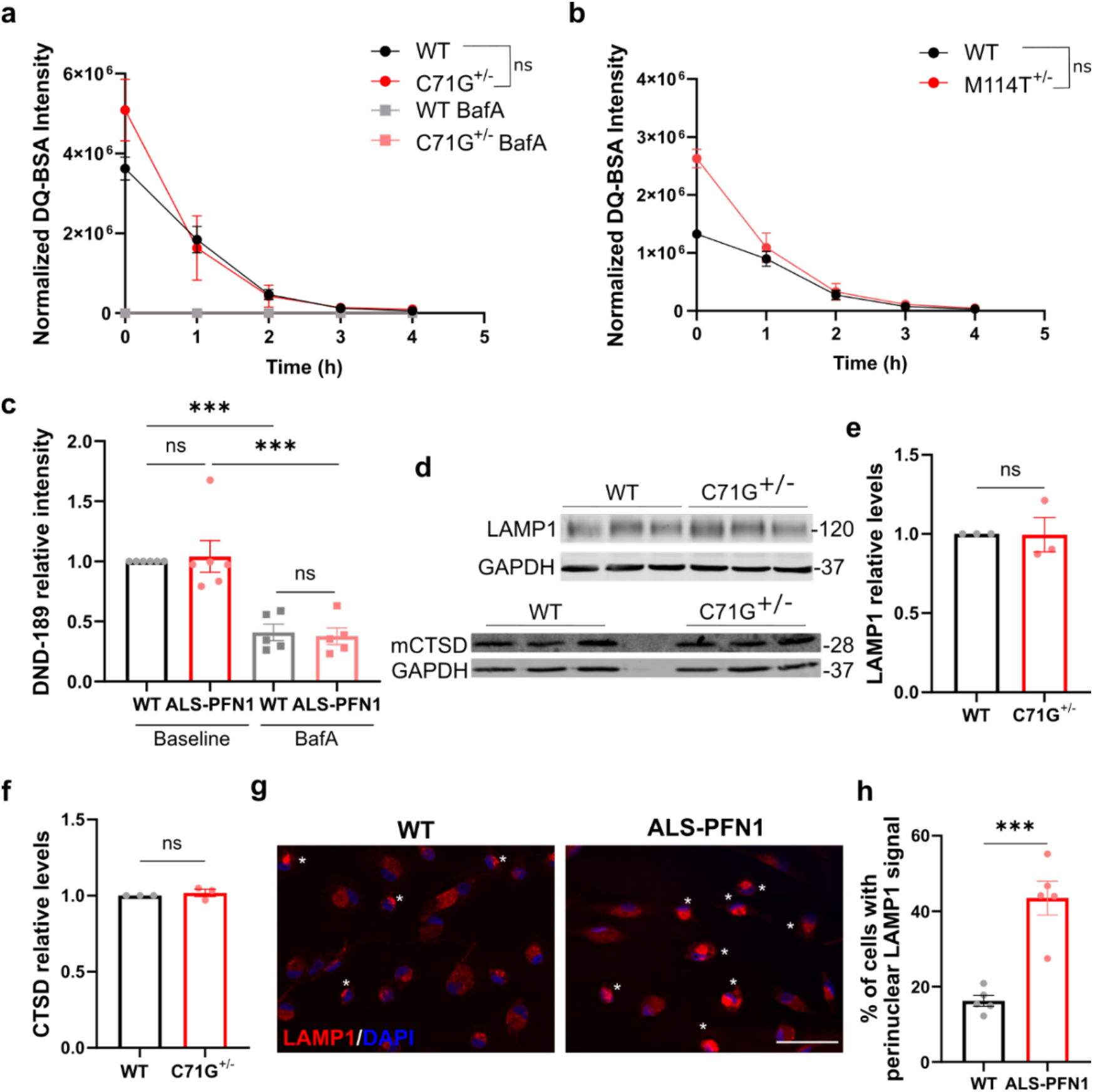
Lysosomal perinuclear accumulation occurs in ALS-PFN1 iMGs. **a,b,** De-quenched DQ-BSA fluorescence intensity measured over 4h by live cell imaging of PFN1 WT and C71G^+/-^ iMGs (**a,** ns *P*=0.5740, F (1, 4) = 0.3737) and of PFN1 WT and M114T^+/-^ iMGs (**b,** ns *P*=0.0717, F (1, 4) = 5.921). Data were normalized to total cell number. Statistics were obtained by two-way ANOVA for n=3 independent differentiations. **c,** The fluorescence intensity of Lysosensor DND-189 was quantified for WT (n=5-6 for two different lines) and ALS-PFN1 (C71G^+/-^ n=3 and M114T^+/-^ n=2-3) iMGs at baseline or pre-treated with 200 nM Bafilomycin A (BafA; *** *P*=0.0006 for WT untreated vs BafA or *P*=0.0002 for ALS-PFN1 untreated vs BafA; ns *P*=0.9840 for untreated WT vs ALS-PFN1 or *P*=0.9938 for BafA WT vs ALS-PFN1). Data was normalized to WT iMGs for each independent differentiation. Statistics were determined by one-way ANOVA F 1.245 (3, 18) and Tukey’s multiple comparisons test. **d,** Western blot analysis of LAMP1 and the mature form of Cathepsin D (mCTSD) from WT and C71G^+/-^ (n=3 independent differentiations) with GAPDH as a loading control. **e,f,** Quantification of **d** for LAMP1 (**e,** ns *P*=0.9703, t=0.03967, df=4) and mCTSD (**f,** ns *P*=0.4999, t=0.7409, df=4). Levels of the target proteins were first normalized to GAPDH and then to the WT control lane corresponding to the same differentiation. Statistics were determined by unpaired two-tailed t-test. **g,** Representative immunofluorescence images of LAMP1 in PFN1 WT (n=7) and ALS-PFN1 (C71G^+/-^ n=4 and M114T^+/-^ n=3) iMGs. White asterisks indicate cells with perinuclear LAMP1 signal. Scale bar=50 µm. **h,** Percentage of iMGs showing LAMP1 perinuclear localization for WT (n=5 for two iPSC lines) and ALS-PFN1 (C71G^+/-^ n=2 and M114T^+/-^ n=3) iMGs. Statistics were determined by unpaired two-tailed t-test (*** *P*=0.0004, t=5.768, df=8). All graphs show mean ± SEM and individual data points in the bar graphs represent independent differentiations.

### Restoring autophagic flux ameliorates phagocytic dysfunction in ALS-PFN1 iMGs

The experiments above implicate ALS-PFN1 in reducing autophagic flux, which can explain the presence of lysosomal clustering. Further, TBC1D15 (**Figure 3**) is also involved in autophagosome morphogenesis and directly interacts with critical autophagy-related proteins (e.g., ATG8 family proteins)^17^. Given these observations, we turned our attention to upstream processes within the endo-lysosomal pathway. Indeed, more signals of early endosome antigen 1 (EEA1)-positive endosomes, which are normally cleared through the autophagy pathway^38^, were detected in PFN1 C71G^+/-^ iMGs relative to controls by immunofluorescence (**Figure 7a,b**). Using a Western blot analysis, we probed for lipidation of the ATG8 family protein microtubule-associated protein 1A/1B light chain 3B (MAP1LC3B or LC3). Lipidation was assessed by the conversion from LC3I to LC3II, where LC3II binds to autophagosome membranes during autophagy initiation^19^. PFN1 C71G^+/-^ iMGs showed a significant reduction in the ratio of LC3-II to LC3I (LC3II/LC3I), while the levels of total LC3 were similar compared to WT iMGs (**Figure 7c-e**). These results are indicative of reduced autophagosome formation and autophagic flux. The autophagy receptor p62/SQSTM1 (sequestosome 1) was significantly elevated in PFN1 C71G^+/-^ iMGs, also consistent with reduced autophagic flux (**Figure 7f,g**). Addition of rapamycin, a known inducer of autophagic flux^39^, restored p62 levels in PFN1 C71G^+/-^ iMGs to that of WT iMGs (**Figure 7f,g**). Strikingly, administration of rapamycin to iMGs prior to the phagocytosis assay with pHrodo-labeled mouse synaptosomes prevented the accumulation of pHrodo signal in ALS-PFN1 C71G^+/-^ and PFN1 M114T^+/+^ iMGs, restoring the phagocytosis index in these mutant iMGs to that of WT iMGs (**Figure 7h,i**). Therefore, targeting the autophagy pathway ameliorated defective phagocytic processing in ALS-PFN1 iMGs.

**Figure 7.**
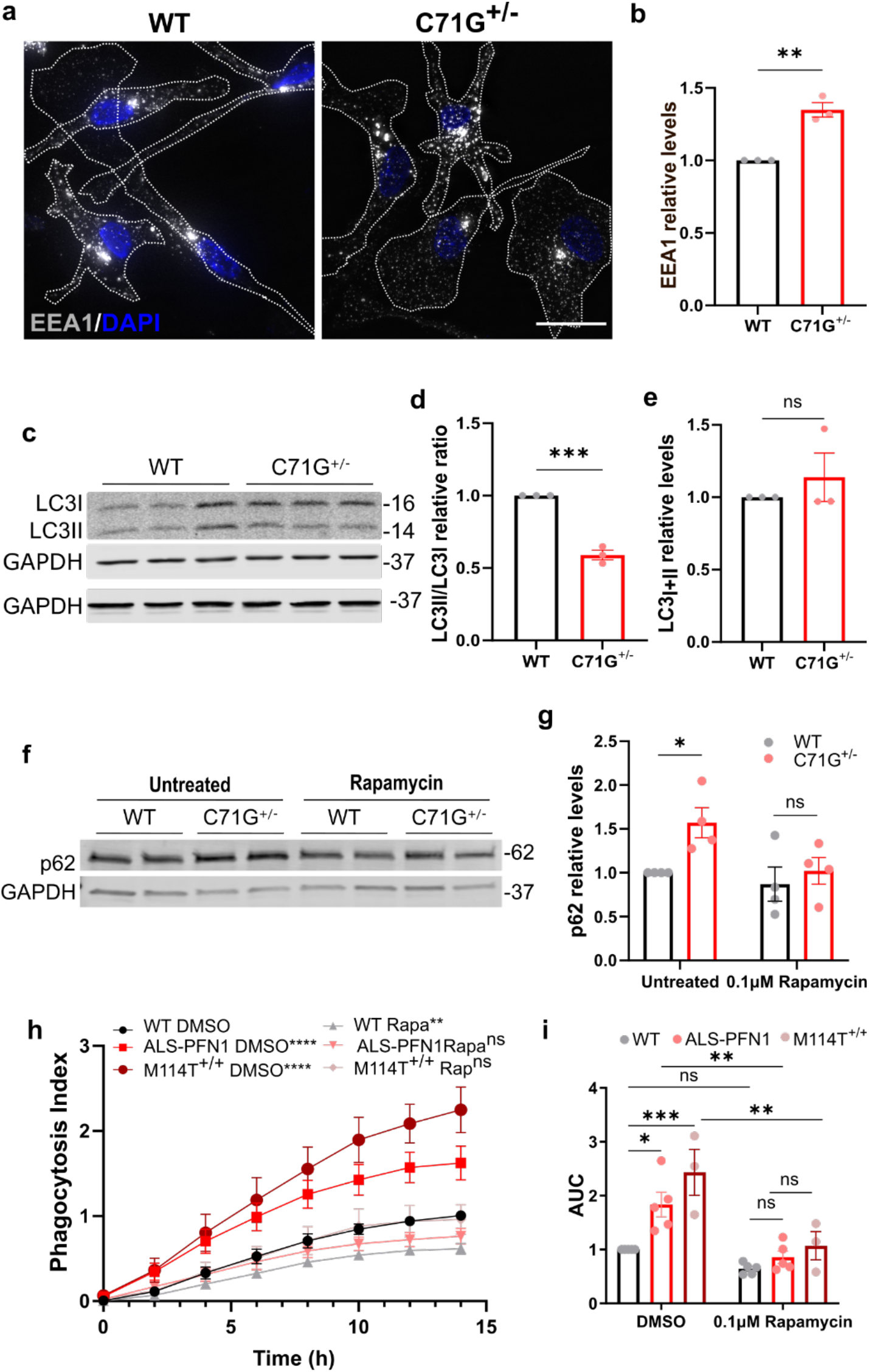
Rapamycin ameliorates deficits in autophagic and phagocytic processes in ALS-PFN1 iMGs. **a,** Representative immunofluorescence images of EEA1. **b,** Quantification of the EAA1 signal intensity measured per cell and normalized to WT iMGs (*\*\*P*=0.0021, t=7.070, df=4). **c,** Representative Western blot analysis of LC3I, LC3II, and GAPDH. **d-e,** Quantification of LC3I and LC3II in **c**. **d,** LC3II/LC3 ratio was determined from **c** and normalized to WT iMGs (*\*\*\*P*=0.0003, t=12.31, df=4). **e,** Total LC3 levels were obtained by summing the intensity values of LC3I and LC3II and normalizing to GAPDH relative to WT iMGs (ns *P*=0.4553, t=0.8259, df=4). **f,** Western blot analysis of p62 for PFN1 WT and C71G^+/-^ iMGs under untreated conditions or pre-treated with 0.1µM rapamycin (n=4 independent differentiations). **g,** Quantification of **f**, with p62 bands normalized to GAPDH and all data reported as relative to untreated WT iMGs. Statistics were performed using two-way ANOVA F (1, 12) = 5.737 and Šídák’s multiple comparisons test (*\*P*=0.0397, ns= 0.7428). **h,i** Live-cell phagocytosis assays using pHrodo-labeled mouse synaptosomes as substrate for WT (n=5 for two different lines), ALS-PFN1 (C71G^+/-^ n=2 and M114T n=3) and M114T^+/+^ (n=3) iMGs pretreated with 0.1µM Rapamycin (Rapa) or DMSO as vehicle. **h,** Quantification of the phagocytosis index (see Methods). Two-way ANOVA F (5, 160) = 70.72 and Tukey’s multiple comparisons test was used for statistical analysis. Comparisons relative to WT iMGs with DMSO are shown (\*\*\*\**P*<0.0001, \*\**P*=0.0015, *P*=0.4638 for ALS-PFN1 with 0.1µM Rapamycin, and *P* >0.9999 for M114T^+/+^ with 0.1µM Rapamycin). **i,** Area under the curve (AUC) determined from **i**. Statistics were determined by two-way ANOVA F (2, 20) = 12.10 and Šídák’s multiple comparisons test for: the DMSO condition (\**P*=0.0326 for WT vs ALS-PFN1 and \*\*\**P*=0.0006 for WT vs M114T^+/+^), the DMSO vs 0.1µM Rapamycin condition per genotype (*P*=0.9152 for WT, \*\**P*=0.0081 for ALS-PFN1, and \*\**P*=0.0038 for M114T^+/+^) and the 0.1µM Rapamycin condition (*P*=0.9993 for WT vs ALS-PFN1, *P*=0.8904 for WT vs M114T^+/+^). Data points in bar graphs represent independent differentiations.

### ALS-linked PFN1 exhibits enhanced binding to PI3P, a lipid required for autophagy

We next sought to explore how PFN1 may influence autophagic flux and considered that PFN1/lipid interactions could be involved given that PFN1 binds phosphoinositides^9, 19, 40^. A particular phosphoinositide of interest, phosphatidylinositol 3-phosphate (PI3P), is generated by vacuolar sorting protein-34 (Vsp-34) and functions as a critical signaling lipid for autophagy^19^. However, potential interactions between PFN1 and PI3P have not been deciphered. To examine whether PFN1 binds PI3P, we first used a differential scanning fluorimetry (DSF) assay that reports on PFN1 melting temperature (T_m_), a measure of protein stability, that we used before to study PFN1 binding with other ligands^41^. In the absence of ligand, the T_m_ of PFN1 M114T is ∼14°C below that of PFN1 WT (**Figure 8a**), consistent with a destabilizing effect of the M114T mutation^41^. Increasing concentrations of PI3P caused a reduction in the T_m_ of both PFN1 WT and M114T (**Figure 8a-c**). Upon addition of 200μM PI3P, PFN1 M114T exhibited a significantly larger decrease in T_m_ (**Δ**T_m_) than PFN1 WT (**Figure 8a-c**). Similar effects on PFN1 T_m_ were observed with the plasma-membrane localized 4,5-bisphosphate (PI(4,5)P2) (**Sup. Fig 8a,b**), a well characterized PFN1 ligand^42^. Intriguingly, the effect of PIP binding to PFN1 had the opposite effect as poly-proline ligands, which stabilized PFN1 (i.e., lead to an increase in T_m_)^25, 41^. The biological reason for PIP-induced destabilization of PFN1 remains to be explored but could be relevant to the role of PIPs in cellular signaling.

We turned next to Nuclear Magnetic Resonance (NMR) spectroscopy to further examine the binding interaction between PFN1 variants and the autophagy-associated PI3P. ^15^N-^1^H heteronuclear single-quantum coherence (HSQC) spectra were collected with either ^15^N-labeled PFN1 WT (**Figure 8d**) or M114T (**Figure 8e**) as a function of increasing PI3P concentration. PFN1 chemical shift perturbations, indicative of PI3P binding, were quantified (**Sup. Fig 8c**) and mapped onto the structure of PFN1 WT in complex with actin and a poly-proline sequence from vasodilator-stimulated phosphoprotein (VASP)^43^; the crystal structure of this ternary complex was used to place chemical shift perturbations arising from PI3P into context with other PFN1 ligand binding sites. PFN1 residues that show major chemical shift perturbations in response to PI3P binding are located at or near poly-proline binding sites and include residues within the N- and C-terminal α-helices (α_1_ and α_4_) and the β-strands behind them (β_1_, β_2_ and β_7_) (**Sup. Fig 8c,d**). Overall, larger chemical shift perturbations were observed in PFN1 WT (**Figure 8f**,) than in PFN1 M114T (**Figure 8g**) upon titration of PI3P (**Sup. Figure 8c**). A global fitting analysis of residues with robust PI3P-induced chemical shift perturbations was performed (**Sup. Fig 9**), resulting in an apparent dissociation constant (*K*_d_) of 1880 ± 190 µM for PFN1 WT and 370 ± 30 µM for PFN1 M114T with respect to PI3P binding. The smaller chemical shift perturbations together with a 5-fold higher binding affinity indicates that PFN1 M114T undergoes smaller conformational changes to accommodate tighter PI3P binding. We also observed precipitation of PFN1 M114T starting at 85 µM PI3P during the titration experiment, whereas PFN1 WT remained soluble with PI3P concentration up to at 1.1 mM. These observations are consistent with reduced PFN1 stability resulting from both the M114T mutation and binding to PI3P (**Figure 8a-c**).

**Figure 8.**
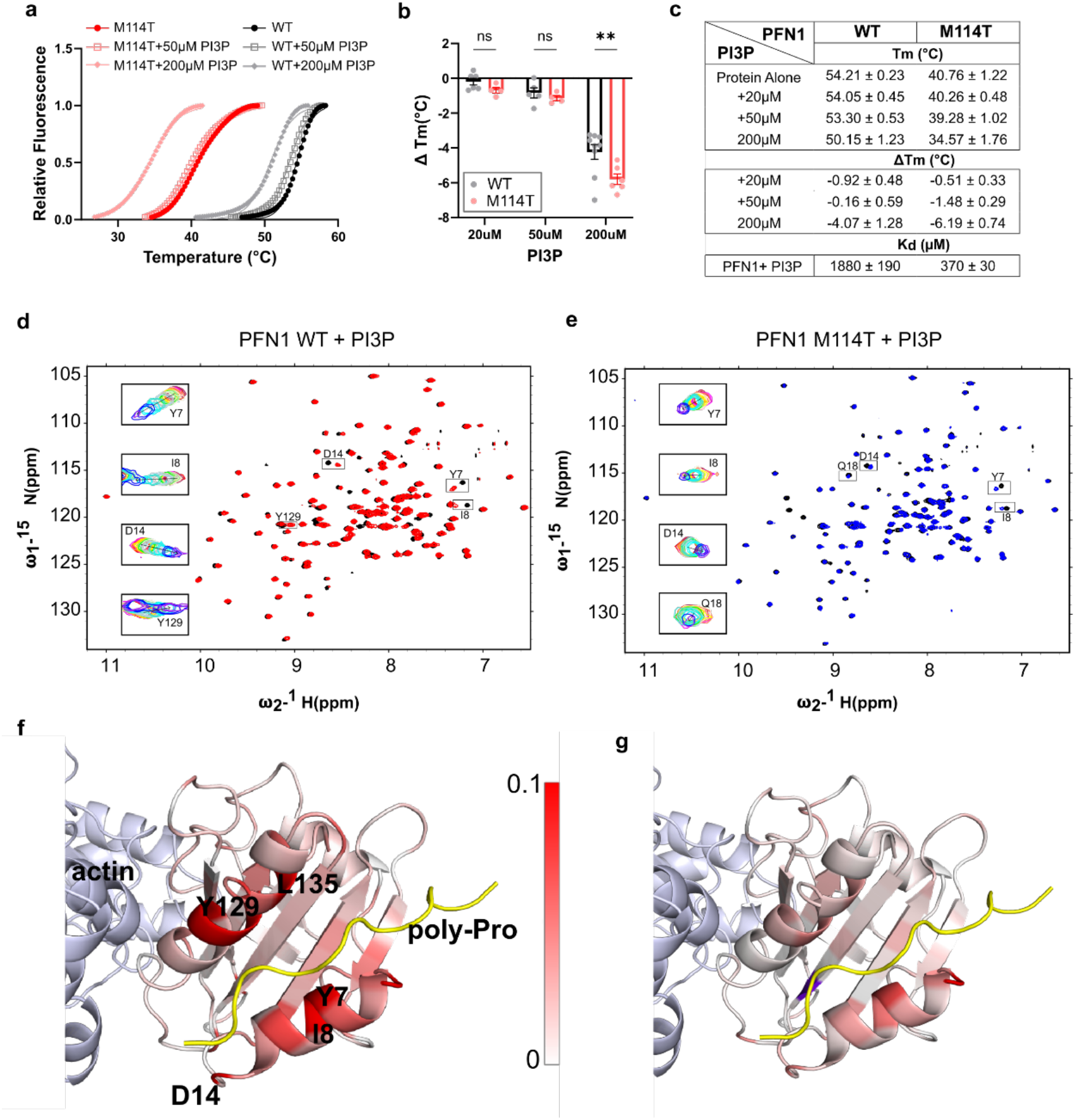
PI3P is a PFN1 ligand with enhanced binding affinity for PFN1 M114T. **a-b,** Differential scanning fluorimetry (DSF) with PFN1 WT and M114T proteins in the presence of PI3P. **a,** Thermal denaturation profiles of PFN1 proteins incubated with the indicated concentration of PI3P measured by SYPRO Orange fluorescence as a function of increasing temperature. An average of two technical replicates is shown and is representative of n=4-9 independent experiments. The curves were fit to the Boltzmann’s sigmoidal function for the determination of an apparent melting temperature (T_m_). **b**, ΔT_m_ reports the difference between the T_m_ at the indicated PI3P concentration and the protein without PI3P. Statistics were determined using two-way ANOVA F (1, 28) = 7.115 and Šídák’s multiple comparisons test (\*\**P*=0.0038 for 200µM and ns *P*=0.735 for 50µM or 0.9326 for 20µM) Bar graphs show mean ± SEM with each data point representing an independent experiment. **c,** Summary of T_m._ and ΔT_m_ obtained from **b** and the dissociation constants from the NMR studies (d-g) for PFN1 WT and M114T in the presence of PI3P. **d-g,** Titration studies of PFN1 with PI3P using NMR. **d,** Overlay of ^15^N-^1^H HSQC spectra of PFN1 free (black) and bound to PI3P (red) for PFN1 WT. **e,** the same as **d** except for PFN1 M114T in the free state (black) and bound to PI3P (blue). **d,e**, Insets show the overlay of spectra collected during the titration of PI3P for select residues. **f,g,** Chemical shift differences between the TROSY 2D ^15^N-^1^H correlation spectra of PFN1 and the PI3P-PFN1 complex are mapped onto the structure of PFN1 in the ternary complex with actin (light blue) and PLP (red), pdb ID 2PAV for PFN1 WT (**f**) and PFN1 M114T PFN1 (**g**). Residues are colored according to the scale bar, with 1 (red) representing PFN1 residues with the greatest chemical shift perturbations upon PI3P binding; chemical shift perturbations are generally for M114T compared to WT. A plot of the chemical shift chemical shift perturbations (ppm) and the secondary structural elements of PFN1 that comprise the three dimension structure in **f,g** are included in **Supplemental** Figure 8.

## DISCUSSION

The mechanism by which mutations in PFN1 cause ALS is unknown. Prior reports with rodent models showed that ALS-linked mutations in PFN1 induce motor neuron degeneration^44, 45^. However, these studies were primarily carried out under conditions of PFN1 overexpression, and none addressed a potential effect of ALS-PFN1 on microglia. While the core functions of microglia are needed to develop and maintain a healthy CNS^1^, microglia dysfunction contributes to neuron degeneration through non-cell autonomous mechanisms^2^. In support of this notion, motor neuron degeneration is modulated by ALS gene expression in microglia^46^. Studies with iPSC-derived microglia cells are providing insights into how disease-linked genes alter the intrinsic properties of human microglia^47, 48^. Though it is widely recognized that the properties of microglia differ between *in vitro* and *in vivo* environments^49^, emerging evidence supports that iMGs are a relevant model for investigating the effects of disease-linked genes on canonical microglia functions. Herein, immunofluorescence, transcriptomics, LPS and phagocytosis studies demonstrate a commitment of ALS-PFN1 and WT iMGs toward a microglia cell fate. Our experiments also revealed that mutant PFN1 disrupts phagocytic processing, a core microglia function. Parallel studies in knock-in PFN1 C71G^+/-^ mice intracranially injected with dead neurons recapitulated a defect in phagocytosis, demonstrating consistency between disease-relevant phenotypes in iMGs and *in vivo*.

Our longitudinal and immunofluorescence analyses showed deficient processing of phagocytosed material within ALS-PFN1 iMGs relative to controls. Evidence from multiple lysosomal functional assays suggest that defective phagocytic processing in ALS-PFN1 iMGs is not caused by reduced lysosomal degradative capacity. Rather, the accumulation of phagocytosed material likely stems from inefficient vesicular processing into and/or through the endo-lysosomal pathway. Intriguingly, perinuclear accumulation of lysosomes was observed in ALS-PFN1 iMGs, a condition that has been observed in other neurological disorders^50^. Lysosome positioning is dependent on cytoskeletal dynamics, which are perturbed by ALS-linked mutations in PFN1^25, 51^. Although ALS-PFN1 iMGs did not exhibit deficiencies in F-actin levels or in the uptake of synaptosomes, both of which require actin polymerization, we cannot exclude the possibility that subtle alterations in cytoskeletal dynamics contribute to inefficient phagocytic processing in ALS-PFN1 iMGs.

Lysosome positioning is also tightly linked to autophagic flux^37, 50, 52^. Thus, perinuclear clustering of lysosomes may signify defects in the autophagy pathway of ALS-PFN1 iMGs. Indeed, ALS-PFN1 iMGs exhibited reduced conversion of LC3I to LC3II, a process that is required for autophagosome formation^19^. Consistent with a reduction in autophagic flux, enhanced levels of the autophagy receptor p62 and EEA1 were detected in ALS-PFN1 iMGs. Genes and proteins associated with the autophagy pathway were also differentially expressed between PFN1 iMGs and control iMGs. Notably, TBC1D15 was upregulated at the transcript and protein levels in ALS-PFN1 iMGs. TBC1D15 is involved in autophagosome morphogenesis during selective autophagy and was shown to interact directly with ATG8 family proteins, including microtubule-associated protein-1 light chain 3 (MAP1LC3) and GABAA receptor-associated protein (GABARAP) orthologues^17, 53^. PFN1 may also serve an important role in autophagy, as it co-immunoprecipitated with multiple ATG8 family proteins^53^ and with Beclin-1^40^, a critical component of the class III phosphatidylinositol 3-kinase (PI3KC3) complex I that forms during autophagy induction^19^. Despite lower steady-state levels of PFN1 C71G, we observed enhanced colocalization of PFN1 with TBC1D15 in ALS-PFN1 iMGs. Further, PFN1 knockdown in HMC3 cells resulted in reduced expression of TBC1D15. These observations implicate an interaction between PFN1 and TBC1D15, potentially within degradative vesicles associated with the autophagy pathway. Upregulation of TBC1D15 may therefore represent a compensatory mechanism to overcome deficiencies in autophagy pathway within ALS-PFN1 iMGs. In addition to Beclin 1, PI3KC3 complex I contains vacuolar sorting protein-34 (Vsp-34), a class III phosphoinositide 3-kinase that catalyzes the production of PI3P^19^, which also interacts with TBC1D15^54^. Here, we identified PI3P as a ligand of PFN1 and found that mutant PFN1 exhibits enhanced binding affinity for PI3P compared to PFN1 WT. Tighter binding between PFN1 M114T and PI3P also resulted in enhanced instability of mutant PFN1, suggesting the functionality of the PFN1/PI3P complex could be compromised in cells. Collectively, the outcomes of our study suggest a model whereby ALS-linked mutations perturb a physiological function of PFN1 in the autophagy pathway, potentially through altered interactions with TBC1D15 and/or PI3P, culminating in impaired autophagic flux.

Our studies with Rapamycin, a potent inhibitor of the mammalian target of rapamycin (mTOR) that induces autophagic flux A, provide a link between autophagic flux and phagocytic processing in ALS-PFN1 iMGs^39^. Treating ALS-PFN1 iMGs with Rapamycin restored p62 and phagocytic processing of human synaptosomes to the same levels as control iMGs. This could be explained by a convergence of the autophagy pathway with phagocytosis through a process called LC3-associated phagocytosis (LAP)^19^. During LAP, LC3 is recruited to phagosomes through a mechanism that requires Beclin-1 and Vsp-34 activity^19^. LAP has been shown to facilitate clearance of pathogens and apoptotic neurons, although less is known regarding the role of LAP in phagocytic degradation of synaptosomes. Aside from LAP, impaired autophagic flux within ALS-PFN1 iMGs could dysregulate other components of the endo-lysosomal pathway, leading to congestion of vesicular degradation in these cells. To date, we have not been able to modulate gene expression in iMGs, as viral transduction was highly toxic to these cells. Recent advances with adeno-associated viruses that induce robust transgene expression in microglia with minimal side-effects may allow for viral transduction of iMGs^55^, thus facilitating future studies aimed at manipulating endo-lysosomal and autophagy pathway components in ALS-linked PFN1 iMGs.

The outcomes of our studies also implicate mutant PFN1 in lipid dysmetabolism. For example, terms related to lipid oxidation and transport were enriched in the iMG proteomics analysis, and elevated lipid droplet content was detected within ALS-PFN1 iMGs. Proper lipid metabolism is important for core microglial functions such as phagocytosis and cytokine secretion^56^, whereas lipid dysmetabolism in microglia is linked to aging and neurodegeneration^29, 56^. PFN1 appears to associate with lipid droplets, which could occur through direct binding of PFN1 with the phospholipid outer shell and thus represent a role for PFN1 in lipid droplet processing. An alternative, though not mutually exclusive, possibility is that impaired autophagic flux results in lipid droplet accumulation. Indeed, lipid droplets are processed by a form of selective autophagy called lipophagy that relies on autophagosome-mediated degradation of lipids^57^. Microglia/macrophages containing excessive lipid droplets exhibit compromised phagosome maturation and enhanced cytokine release^29, 58^. Herein, we did not detect differences in cytokine release as a function of PFN1 genotype in iMGs, even when iMGs were challenged with LPS. It is possible that the inflammatory response of microglia expressing ALS-linked PFN1 is altered by conditions not tested herein, such as *in vivo*, as a function of age and/or due to chronic accumulation of debris. Expanding our iMG studies into a co-culture format with neurons, as well as further investigations with the novel PFN1 C71G^+/-^ mouse model presented herein, may reveal additional phenotypes induced by ALS-PFN1^6^. In the context of lipid metabolism, glia were shown to buffer lipids from neighboring neurons under conditions of neuronal stress *in vivo*^59^. Further investigation into whether lipid dysmetabolism in glia precludes this neuroprotective function is warranted, as this may represent a mechanism by which microglia modulate neurodegeneration.

Our results have implications for ALS and related neurodegenerative disorders. In the context of ALS-PFN1, the outcomes of our study raise the possibility that microglial dysfunction caused by mutant PFN1 could directly affect the CNS microenvironment, and by extension the health and viability of motor neurons. Intrinsic microglia dysfunction may be more prevalent in ALS than previously thought, as other ALS-associated genes including *C9ORF72* are abundantly expressed in microglia. Further, multiple ALS-linked proteins, including tank-binding kinase (TBK1), optineurin (OPTN), ubiquilin 2 (UBQLN2), C9ORF72, and p62 are expressed in microglia and are directly involved with autophagy^60^. While there has been an emphasis on the autophagy pathway in neuronal subtypes across different neurodegenerative diseases, targeting canonical and non-canonical autophagy pathways in microglia may also be a viable therapeutic strategy for these disorders^61^.

## METHODS

### CRISPR engineering of PFN1 mutant iPSC lines

The PFN1 C71G and M114T lines and their respective isogenic control lines were generated from the euploid (46; XY) KOLF2.1J iPSC line ^21^ at The Jackson Laboratory for Genomic Medicine (Farmington, CT). The PFN1 C71G single-nucleotide variant T > G was generated using a guide RNA that overlaps codon 211 of PFN1 while the PFN1 M114T single-nucleotide variant T > C was generated with a guide RNA that overlaps codon 341 as listed below. A 100-nt single-stranded oligonucleotide donor template was used for both mutations as described before with minor modifications^62^. The Amaxa 4D nucleofection system (Lonza) was used as follows: pre-assembled Cas9 RNP (2 µg IDT HiFi V3 Cas9 and 1.6 µg Synthego synthetic modified single guide RNA) and 40 pmol single-stranded oligonucleotide (IDT; HDR Donor oligo) were added to 20 µL P3 buffer containing 1.6 × 10^5^ Accutase-dissociated iPSCs and the nucleofection procedure was carried out in a 16-well Amaxa 4D cuvette (Lonza; Primary Cell P3, pulse code CA137). Cells were cultured for 3 days under cold-shock conditions (32 °C/5% CO_2_) and in the presence of RevitaCell (ThermoFisher) and HDR enhancer (1 µM Alt-R HDR Enhancer v2, IDT) for the first 24 h. Once the cells were confluent, iPSCs were dissociated and plated at low density to obtain single-cell-derived clones. Clones were screened by Sanger sequencing of PCR amplicons to identify the desired homology-directed repair events in PFN1 using the primers below.

For the PFN1 C71G variant:

Cas9 guide RNA: CCCGGATCACCGAACATTTC

Repair oligo: TCTTGGTACGAAGATCCATGCTAAATTCCCCATCCTGCAGCAGTGAGTCCCGGATCACCGAACCTTTCTGGCCCCCAAGTGTCAGCCCATTCACGTAAAA

PCR primer (forward): 5′-GAATCTTGGTGCACTGACTAACTTG and PCR primer (reverse): 5′-AAGAACTCAAACGATGAACTCGATG

Sanger sequencing primer: 5′-ACTAACTTGATGGGCGCTTG

For M114T variant:

Cas9 guide RNA: CCAGCGCTAGTCCTGCTGAT

Repair oligo: GGTGGGAGGCCATTTCATAACATTTCTTGTTGATCAAACCACCGTGGACACCTTCTTTGCCCGTCAGCAGGACTAGCGCTGGAGGAGGAGGAAAGAGAAA

PCR primer (forward): 5′-CACATGAATCAACAGTCTATGAGCC and PCR primer (reverse): 5′-CACACAGCACCTTGTTAGTAGAATC

Sanger sequencing primer: 5′-GTTGGGAGAGATGAGGTTGG

### Generation of human iPSC-derived microglia-like cells

The microglia differentiation method used in this study has been previously described by us in detail^8^. In summary, iPSC colonies were maintained in mTeSR Plus media (Stem Cell Technology, 100-0276) and passaged every 4-6 days with 0.5mM ethylenediaminetetraacetic acid (EDTA, Invitrogen 15575020) in Dulbecco’s Phosphate-Buffered Saline (DPBS; Corning, 21-031-CV). Six-well plates coated for at least 2h with 10µg/mL of Cell Adhere Laminin 521 (Stem Cell Technology, 200-0117) diluted in DPBS containing calcium and magnesium (Corning, 21-030-CV) were used to increase the yield of downstream progenitors as described ^63^. Before differentiation, expression of the pluripotency markers octamer-binding transcription factor 4 (OCT4) and SOX2 were confirmed in all iPSC lines by immunofluorescence staining (**Sup Fig1**). Induced PSCs were differentiated within five passages into microglia-like cells (iMGs).

On day 0, iPSCs were differentiated into embryoid bodies (EBs)^63, 64^. When iPSC colonies were 80% confluent, cells were dissociated using TrypLE reagent (Life Technologies, 12-605-010) for 2 minutes at 37°C, gently detached using a cell lifter and collected in DPBS. The cells were pelleted by centrifugation at 500 x g for 1 min and resuspended in EB media (**Table 1**) to induce a mesoderm lineage. Cells were seeded at a density of 10^4^ cells/well into low adherence 96-well plates (Corning 7007) and centrifuged at 125 x g for 3 min. Half the volume of media was replaced with fresh media on day 2.

**Table 1.**
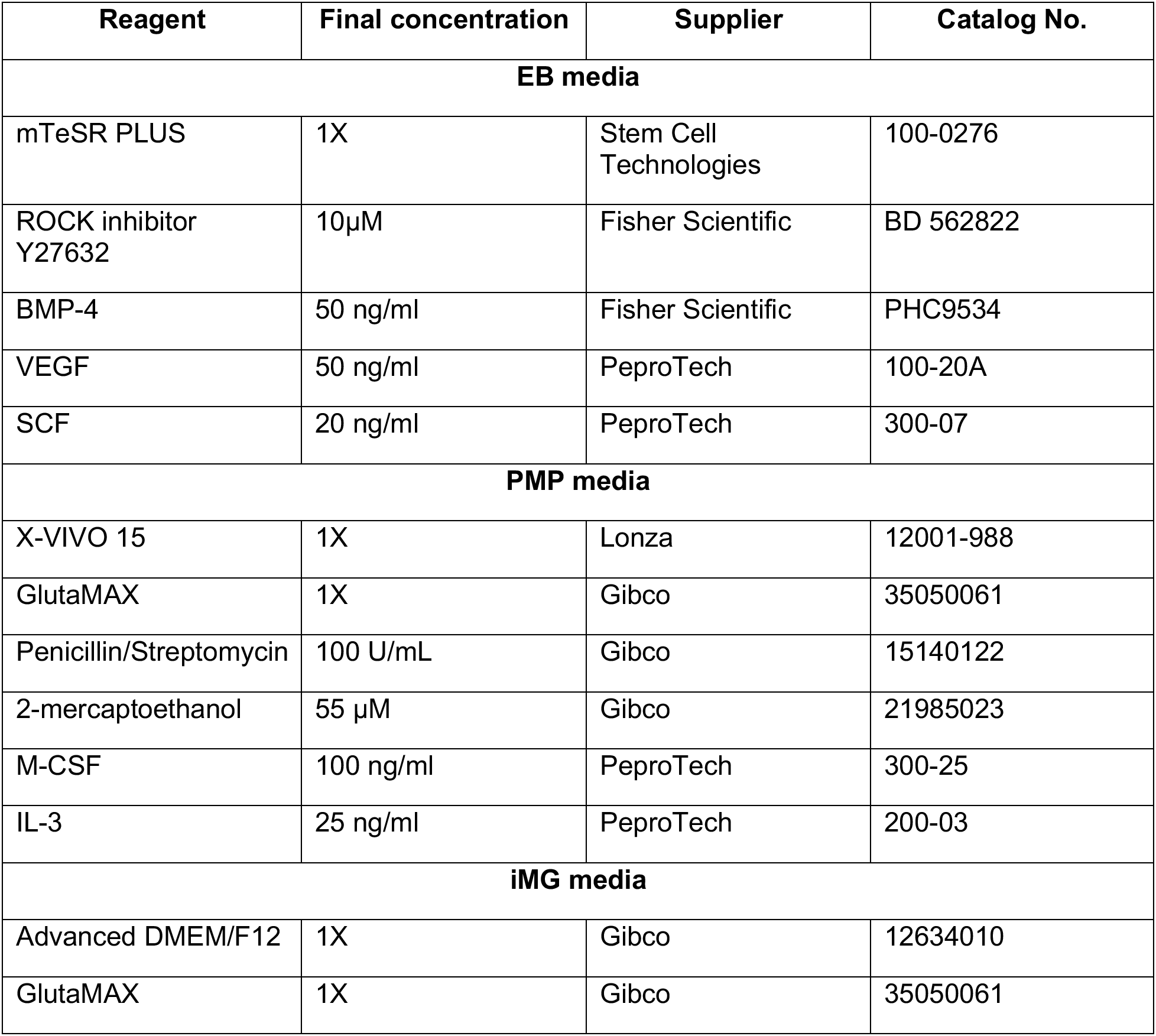
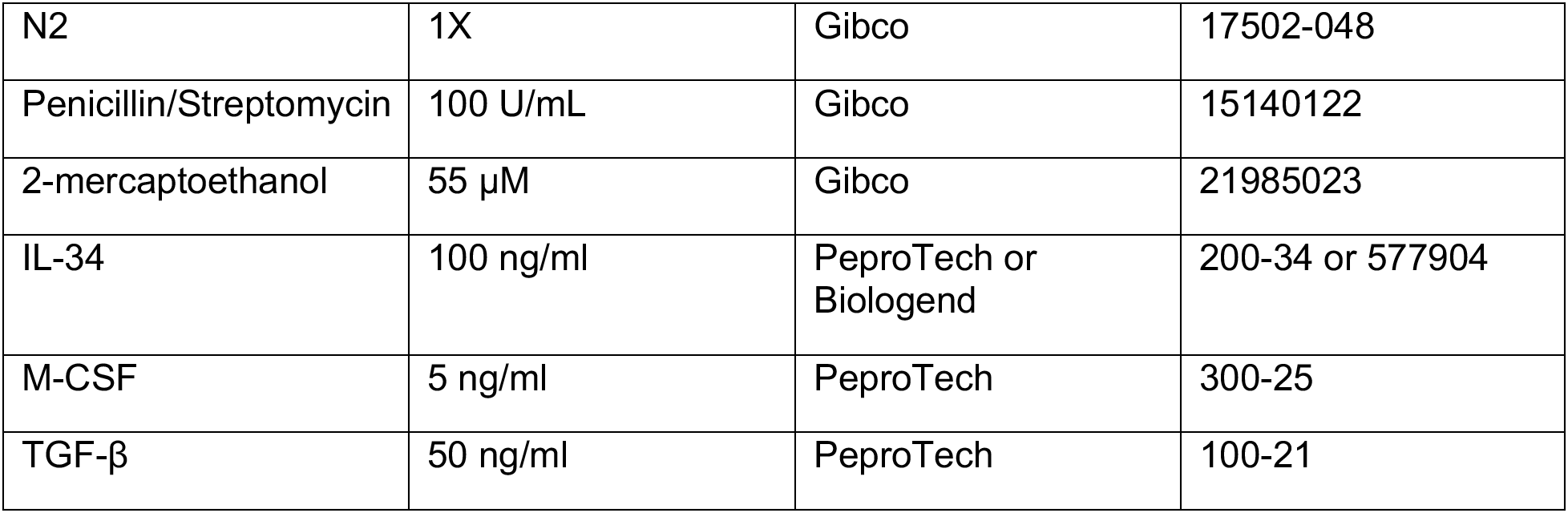
Media and reagents used to generate iPSC-derived microglia-like cells.

On day 4, myeloid differentiation was induced as previously described^5, 6^ with the following exceptions. EBs were plated in 6-well plates pre-coated for at least 1h with Matrigel (Corning, CB-40230) and EBs were diluted in KnockOut Dulbecco modified Eagles media (DMEM)/F-12 (Gibco, 12660012) to improve their attachment onto the plate^63^. EBs were maintained with PMP media (**Table 1**) and a full media change was performed every 5-7 days. After 28 days, primitive macrophage precursors (PMPs) were harvested weekly for up to 3 months by completely removing the PMP media (containing the cells) and then replacing fresh PMP media back into the well. Harvested cells were plated on Primaria-treated cell culture plates (Corning, 353846, 353847, 353872) at a density of ∼10^5^/cm^2^ and terminally differentiated into iMGs for 10-12 days by addition of iMG media (**Table 1**) that contains 100 ng/ml IL-34^5^, 5 ng/ml M-CSF^65^, and 50 ng/ml TGF-β ^66^. Half the volume of media was replaced with fresh media every 3-4 days. All iMG analyses were performed at 10-12 days of microglia differentiation.

### Gene expression analysis by real-time quantitative PCR (qPCR)

Total RNA was isolated using Trizol (Invitrogen,15596026) and treated with DNase I (Invitrogen, 18068015) following the manufacturer’s protocols. RNA purity and concentration were determined from absorbance (A) measurements using a Nanodrop device (ND-1000). Samples with A260nm/A280nm between 1.8 and 2.2 were considered pure and used for downstream applications. Complementary DNA (cDNA) was synthesized using iScript Reverse Transcription Supermix (Bio-Rad, 9170-8840) according to the manufacturer’s instructions and qPCR was performed using 5x iTaq Universal SYBR Green Supermix (Biorad, 172-5120) with the primer sets listed below. The qPCR experiments were conducted at 95 °C for 2 min, 40 cycles at 95 °C for 5s and 60 °C for 30s, with a melt-curve from 65 to 95 °C (in 0.5 °C increments per 5 s) using CFX384 Touch Real-Time PCR Detection System (Bio-Rad, 1845384). Quantification of mRNA was determined using the comparative cycles to threshold CT (ΔΔCT) method, normalizing to *GAPDH* as internal standard and then normalizing to values obtained from WT iMGs for each independent differentiation. Samples were run in triplicate from three independent differentiations or iPSC passages. Averages of technical triplicates per experiment are reported.

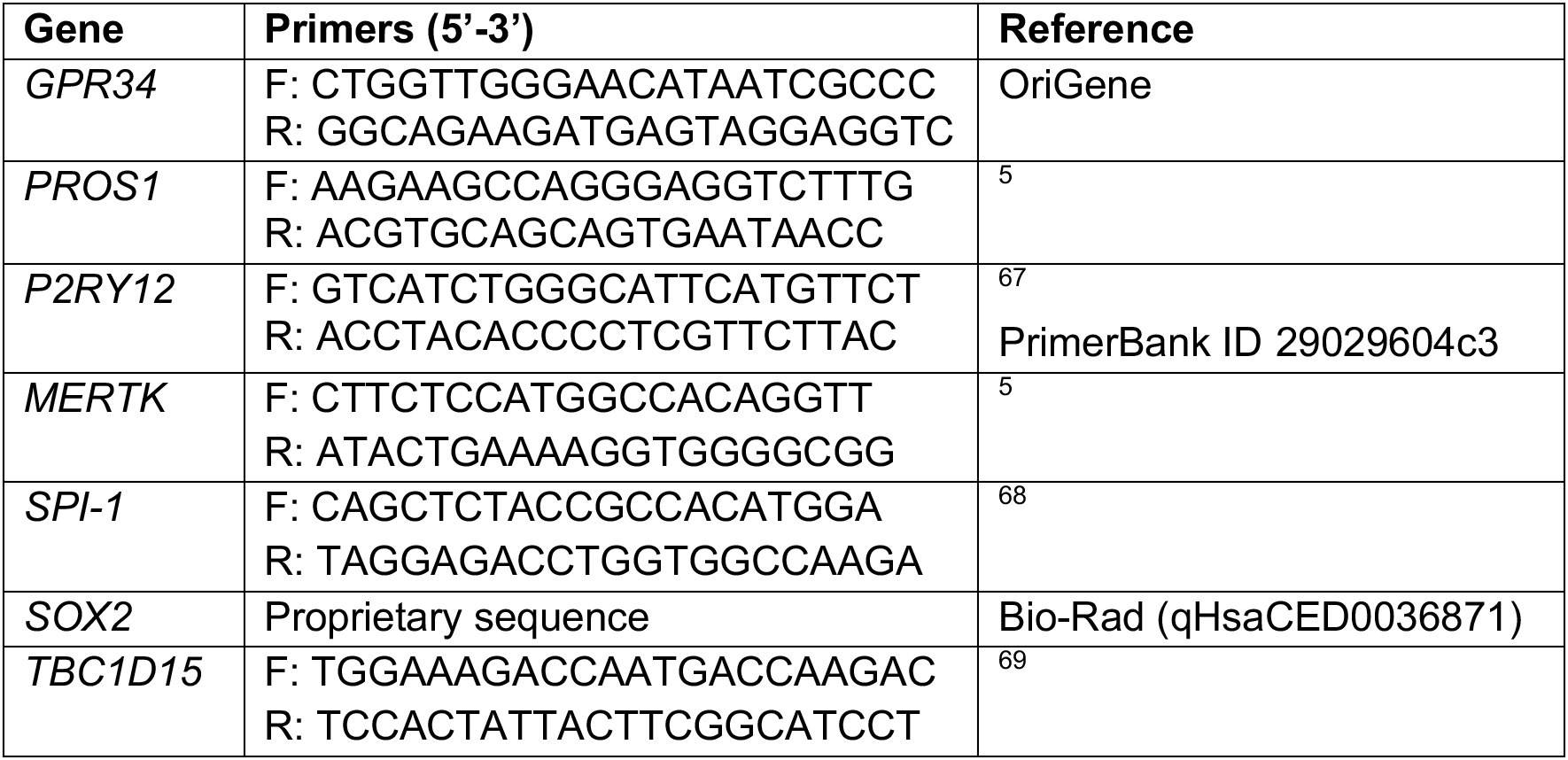

### Transcriptome analysis by RNA sequencing (RNAseq)

Seven independent differentiations from WT iMGs (two different iPSC lines), four independent differentiations from C71G^+/-^ iMGs and three independent differentiations from M114T^+/-^ and M114T^+/+^ iMGs were lysed with Trizol and total RNA was isolated with Direct-zol RNA Microprep Kit (Zymo Research, R2060) according to the manufacturer’s instructions. RNA library preparation and sequencing was conducted by Novogene Co., LTD (Beijing, China) with a read length of 150 base pairs.

Sample sequences were checked for overall quality as well as possible adapter contamination using FASTQC and multiqc tools^70, 71^. FASTQ files were mapped to GRCh38 genome using STAR aligner (2.7.0a) in two-pass mode with a splice aware option^72^. In particular, the options, – outSAMtype BAM SortedByCoordinate was used to produce sorted bam and –sdjbOverhang 100 for optimal splice junction overhang length. Read counts were computed for each transcript based on GENCODE version 33 annotation using HTSeq tool in strand-specific mode^73^. Transcripts with low count reads were filtered from the dataset by retaining only those with a sum of at least 20 reads in all samples.

Principal component analysis (PCA) was performed including previously reported data sets downloaded from Gene Expression Omnibus with accession numbers GSE110952 from Brownjohn et al., 2018^5^ and GSE89189 from Abud et al., 2017^4^. All data sets were batch-effect corrected using ComBat-seq^74^ and normalized using the mean of ratios method in DESeq2^75^. The EdgeR library^76^ was used for expression-based filtering using the filterbyExpr function and counts per million (CPM) were calculated. PCA was performed with CPM values using the prcomp function and the plotly package for 3D visualization^77^ in R.

Differential gene expression analysis was performed using read counts with the DESeq2 package. The Wald test was used to obtain *P*-adjusted values and log2 (fold change) between the WT group (n=7 WT samples from two iPSC lines) and the ALS group (n=4 C71G^+/-^ and n=3 M114T^+/-^ samples)^75^. Genes with a *P*-adjusted value<0.05 were considered differentially expressed.

### Lipopolysaccharide (LPS) stimulation and cytokine analysis

Induced microglia-like cells were treated for 6 and 24h with 100ng/ml of LPS from *Escherichia coli* (*E. coli*) O55:B5 (Sigma L2880) that was diluted first in DPBS and then added to the iMG media. After each time point, the supernatant was collected, centrifuged at 14,000 x g for 10 min and stored at -80°C. Samples were thawed and the levels of the cytokines were detected with the Human ELISA MAX™ Deluxe Set containing IL-6 (430504), IL-10 (430604), CCL5 (440804), and TNF-α (430204) from BioLegend according to the manufacturer’s protocol. Each condition was measured in duplicate and the average of three independent differentiations is reported herein.

### Quantitative proteomics

One million cells from three independent differentiations for WT and C71G^+/-^ iMGs were lysed with RIPA buffer (Boston BioProducts, BP-115-500) supplemented with protease inhibitors (Roche, 11836170001) on ice for 15 min. Lysates were cleared through centrifugation (14,000 g for 10 min at 4°C) and total protein content was quantified with a Bicinchoninic acid assay (BCA) Assay (Thermo Scientific Pierce, 23227) according to the manufacturer’s instructions. For each sample, 50 μg of lysate was diluted to a final concentration of 1 μg/μL with 100 mM triethyl ammonium bicarbonate for tandem mass tag (TMT) labeling using an amine-reactive TMT Isobaric Mass Tagging 6-plex Kit (Thermo Fischer, 90061) as previously described ^78^. LC-MS/MS experiments were performed in triplicate using an Orbitrap Fusion Lumos Tribid mass spectrometer in OTOT (orbitrap/orbitrap) ddMS2 mode (Thermo Scientific) equipped with a nanoACQUITY UltraPerformance LC (UPLC; Waters, Milford, MA, USA). The peptides were loaded and trapped for 4 minutes with 5% acetonitrile (0.1% formic acid) at 4.0 µL/min onto a 100 µm I.D. fused-silica precolumn (kasil frit) packed with 2 cm of 5 µm (200Å) Magic C18AQ (Bruker-Michrom, Auburn, CA). Next, the peptides were eluted and separated at 300 nL/min by an in-house made 75 µm I.D fused silica analytical column (gravity-pulled tip) packed with 25 cm of 3 µm (100Å) Magic C18AQ (Bruker-Michrom, Auburn, CA). The UPLC method used for the separation and elution was a 145 minute gradient starting from 10% of acetonitrile (0.1% formic acid) and 90% of water (0.1% formic acid). The MS data acquisition was performed in positive electrospray ionization mode (ESI+), within the mass range of 375-1500 Da with the orbitrap resolution of 120000 with a maximum injection time of 50 milliseconds. Data Dependent Acquisition (ddMS2) was carried out with a 1.2 Da isolation window, with a resolution of 30000 at m/z 200. The maximum injection time for the ddMS2 was set to 110 milliseconds with the customed AGC target with a 38% of HCD collision energy.

Raw data files were peak processed with Proteome Discoverer 2.1.1.21 (Thermo Fisher Scientific Inc.) using a Mascot Search Engine (Matrix Science Ltd, Server version 2.6.2) against the Human (Swissprot, V42) FASTA file (downloaded 04/09/2019). Search parameters included Trypsin enzyme, and variable modifications of oxidized methionine (+16 on M), pyroglutamic acid for glutamine (-17 on peptide N-Terminal) and N-terminal acetylation (+42). Fixed modifications were set for the carbamidomethylation on cysteine (+57 on C) and TMT 6-plex (+229 on peptide N-Terminal and Lysine). Assignments were made using a 10-ppm mass tolerance for the precursor and 0.05 Da mass tolerance for the fragments. The FDR (1%) analysis was carried out by Scaffold (version 4.4.4, Proteome Software, Inc.) and Q+ quantitative analysis was carried out for fold change analysis. PFN1 C71G^+/-^ and WT were compared with a T-test followed by a Benjamini-Hochberg test for multiple-test correction. Proteins with a *P*-value<0.00160 were considered significantly different between genotypes.

### Functional enrichment analysis

Pathway and process enrichment analysis was conducted with differentially expressed proteins (*P*-value<0.00160) using Metascape (https://metascape.org) with the following ontology sources: KEGG Pathway, GO Biological Processes, Reactome Gene Sets, Canonical Pathways, Cell Type Signatures, CORUM, TRRUST, DisGeNET, PaGenBase, Transcription Factor Targets, WikiPathways, PANTHER Pathway, and COVID. All genes in the human genome were used as enrichment background and the default settings were selected to perform the analysis. Functional terms with a *P*-value < 0.01, a minimum count of 3, and an enrichment factor >1.5 were collected and grouped into clusters based on their membership similarities. Clusters are represented by the most statistically significant term within a cluster ^79^. Additionally, enriched gene ontology terms were annotated with Enrichr (https://maayanlab.cloud/Enrichr/) using the libraries: GO_Cellular_Component_2021, GO_Molecular_Function_2021, and GO_Biological_Process_2021 ^80^.

### Western blot analysis

Cell lysates were prepared as described under “Quantitative proteomics”. Western blotting was performed as previously described ^62^ except that for the analysis of the autophagy-related proteins, microtubule-associated protein 1A/1B-light chain 3 (LC3) and sequestosome-1 (SQSTM1 /p62), the transfer was carried using a Trans-Blot Turbo Transfer System (Bio-Rad, 1704150) set with the high molecular weight protocol (1.3A, 25V, 10min), with the Trans-Blot Turbo RTA Mini Transfer Kit (Bio-Rad, 1704272) and 0.45 µm Immobilon-FL transfer membranes (Fisher Scientific, IPFL00010). Primary antibodies were used as follows: 1:1000 for rabbit anti-PFN1 (Sigma-Aldrich, P7749), rabbit anti-CTSD (Cell Signaling Technology, 2284S), rabbit anti-RAB7 (Cell Signaling Technology, 9367T), rabbit anti-LC3 (Cell Signaling Technology, 3868), guinea pig anti-p62 (PROGEN, GP62-C); 1:500 for rabbit anti-TBC1D15 (Sigma-Aldrich, HPA013388), and 1:10000 for mouse anti-GAPDH (Sigma-Aldrich, G8795) or rabbit anti-GAPDH (Sigma-Aldrich, G9545). Membranes were incubated with LI-COR secondary antibodies (Li-Cor Biosciences) at 1:10,000 for 1h as described ^81^. Western blots were visualized using the Li-Cor Odyssey system (ODY-2215) and densitometric analysis was performed using Image Studio (Li-Cor, v 3.1.4). Data were obtained from at least three independent iMG differentiations or individual HMC3 experiments.

### Immunofluorescence, imaging acquisition, and quantification

Immunostaining for fluorescence microscopy was performed as previously described^82^. Briefly, cells grown on glass coverslips (Carolina Biological Supply, 633029) were fixed with 4% paraformaldehyde (Fisher Scientific, AAA1131336) for 15 min and permeabilized with 1% Triton X-100 (Sigma, T9284) for 10 min. Cells were blocked with 50mM ammonium chloride (Cambridge Isotope Laboratories, NLM-467-5), 0.01% bovine serum albumin (BSA; Fisher BioReagents, BP1600), 2% goat serum (Sigma-Aldrich G9023), 0.01% Triton X-100 in PBS for 1h and probed with the antibodies listed below for 1h. After 3-5 washes, cells were incubated with secondary antibodies at 1:2000 for 1h ^81^. For samples in which F-actin levels were analyzed, Alexa Fluor 647-conjugated Phalloidin (Invitrogen, A22287) was added for 45 min following the manufacturer’s recommendations. Cells were stained with DAPI (Sigma-Aldrich, D9542) and coverslips were mounted with ProLong Gold anti-fade reagent (Invitrogen, P36930).

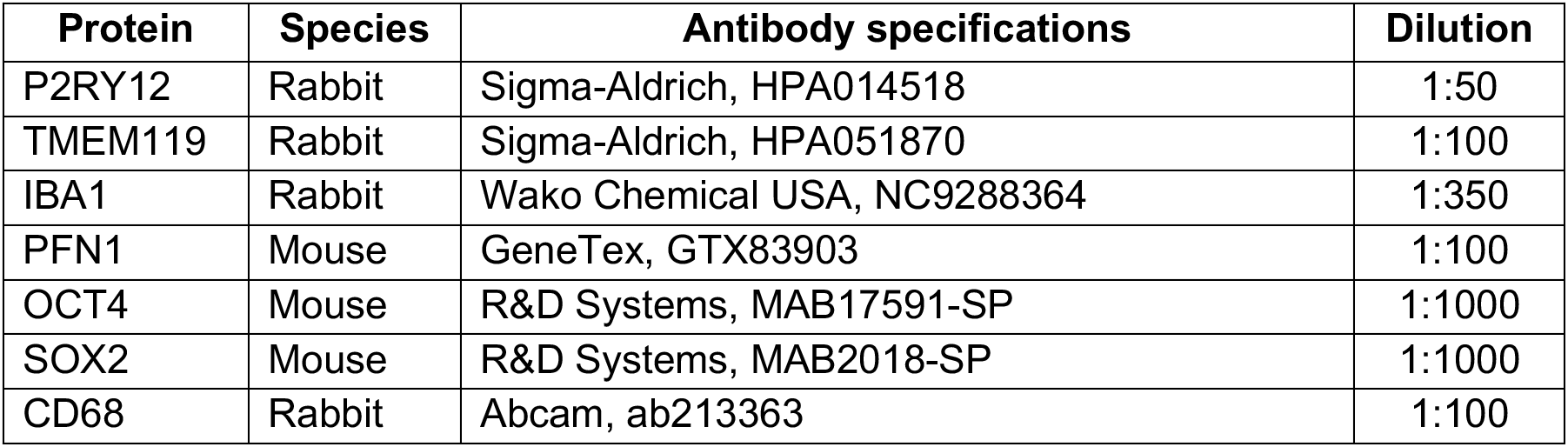
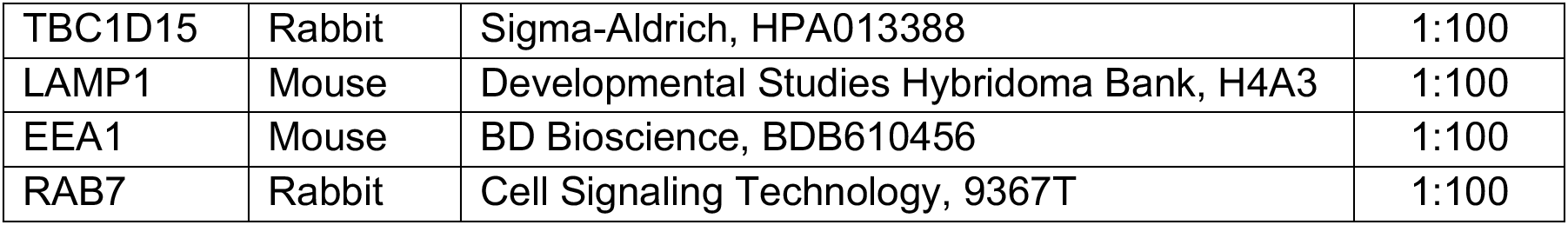

Images were acquired with a 40X air objective, a 63X-oil or a 100X oil-immersion objective on a Leica DMI 6000B inverted fluorescent microscope equipped with a Leica DFC365 FX camera and using AF6000 Leica Software v3.1.0 (Leica Microsystems). Z-stacks were collected with a step size of 0.2 µm and deconvolved using the LAS AF One Software Blind algorithm (10 iterations) from Leica Software v3.1.0. Brightness and contrast were equally adjusted post-acquisition across all images using ImageJ (Fiji, 2.3.051, http://imagej.nih.gov/ij/index.html, USA) to improve target visualization. Images are presented as maximum projections unless otherwise noted.

To calculate the relative area or intensity level of a target protein, individual cells were automatically thresholded using the “Default” method in ImageJ. The “Measure” function was applied to obtain the area and the mean intensity from 60-75 cells across at least three independent differentiations. The experimentalist was blinded to the genotype of the cells. The mean for mutant iMGs was normalized to that of WT iMG corresponding to the same independent differentiation.

Colocalization analyses were performed using JACoP plugin (Fiji, ImageJ) to obtain Pearson’s correlation coefficients and M1 and M2 Mander’s coefficients for individual cells ^83^. Twenty cells were analyzed per condition from three independent microglia differentiations. Cytofluorograms were generated for individual cells to exclude the possibility that bleed through signal or noise are affecting the coefficient values ^83^. Additionally, fluorescent signal intensity plots were obtained for intracellular structures using Leica Application Suite X (LASX) software v3.7.2.

For the analysis of LAMP1 subcellular localization, the number of cells that exhibited LAMP1 signal clustered in the perinuclear area as well as the total number of cells were manually counted for 6-7 fields of view (FOV) per condition using the Cell Counter plugin (Fiji, ImageJ). The experimentalist was blinded to the genotype of the cells. The sum of the number of cells with LAMP1 perinuclear localization was divided to the total number of cells in all the analyzed FOV per condition and multiplied by 100 to obtain the percentage. Images from five WT microglia differentiations derived from two independent iPSC lines, five ALS-PFN1 differentiations derived from M114T^+/-^ iPSCs (n=3) and C71G^+/-^ iPSCs (n=2), and three differentiations derived from M114T^+/+^ iPSCs were used in this analysis. A total of 100-200 iMGs were analyzed per group.

### iPSC-derived lower motor neuron (i^3^ LMN) differentiation

A WTC11 iPSC line with stable insertion of the hNIL inducible transcription factor cassette containing neurogenin-2 (NGN2), islet-1 (ISL1), and LIM homeobox 3 (LHX3) into the CLYBL safe harbor locus was generously provided by Dr. Michael Ward and differentiated as previously described ^84^. Differentiation was induced with DMEM/F12 (Gibco, 11330032) supplemented with N2 supplement, GlutaMAX, nonessential amino acids (NEAA; Gibco, 11140050), 10 μM ROCK inhibitor (Fisher Scientific BD, 562822), 0.2μM compound E (Calbiochem, 565790), and 2 μg/mL doxycycline (Sigma, D9891). After 2 days, the cells were dissociated with Accutase (Corning, 25058CI) and plated on 10-cm cell culture dishes (Corning, 430591) coated with poly-L-ornithine (Sigma, P3655), poly-D-lysine (Sigma, P7405), and 15μg/mL laminin (Gibco, 23017015) at a density of ∼50,000 cells per cm^2^. The cells were maintained with DMEM/F12 containing 10 μM ROCK inhibitor, 0.2μM compound E, 2 μg/mL doxycycline, 40 μM BrdU, and 1μg/mL laminin for another 2 days. For motor neuron maintenance, the neurons were cultured with the B-27 Plus Neuronal Culture System (Gibco, A3653401) supplemented with N2, NEAA, GlutaMAX, and 1μg/mL laminin. Half-media changes were performed every 3-4 days. The cultures were maintained for 28 days before synaptosome purification as described below under “Mouse and human synaptosome purification and labeling”.

### Animals

All mice were maintained in a standard barrier facility at UMass Chan Medical School. The Institutional Animal Care and Use Committee (IACUC) at UMass Chan Medical School approved all experiments involving animals. Standard breeding techniques were used for the generation of wild-type *C57BL/6J* embryos or pups for purification of synaptosomes or isolation of primary cortical neurons, respectively, as described below. *C57BL/6J-Pfn1^em4Lutzy^/J* mice (#030313) were generated by and obtained from the Jackson Laboratory (Bar Harbor, ME). These mice were CRISPR engineered to harbor the allele *Pfn1^C71G, R75R^*. The cohort of mice used in this study was generated by *in vitro* fertilization at the UMass Chan Medical School Transgenic Animal Modeling Core (TAMC; Worcester, MA) using male *C57BL/6J-Pfn1^em4Lutzy^/J* heterozygous mice and female wild-type *C57BL/6J* mice.

### Mouse and human synaptosome purification and labeling

Mouse synaptosomes were purified from wild-type *C57BL/6J* mice postnatal day 18-21. The following steps were performed on ice and under sterile conditions. Cortices were dissected and homogenized in sucrose-HEPES buffer [0.32M sucrose (Fisher Scientific, S5-500), 5mM N-2-hydroxyethyl piperazine-N-2-ethane sulfonic acid (HEPES; Gibco, 15630080), and cOmplete, mini protease inhibitor cocktail tablets (Roche, 11836153001)] using a glass Dounce homogenizer with 50 strokes. The homogenate was centrifuged at 1,200 xg at 4°C for 10 min to clear cell debris. The supernatant was collected and centrifuged at 15,000 xg for 15 min. The resulting pellet was resuspended in 1mL of sucrose-HEPES buffer. A discontinuous gradient containing 0.8M, 1M, and 1.2M sucrose (from top to bottom) diluted each in 5mM HEPES was prepared in 13.2 mL open-top, thin-wall ultraclear tubes (Beckman Coulter, 344059). The collected supernatant was layered over the sucrose gradient and centrifuged at 150,000 xg for 2 h in a swinging bucket rotor at 4°C. Fractions at the interface between the 1.0 and 1.2 M sucrose layers were collected, diluted to 0.32M sucrose by adding 2.5 volumes of 5 mM HEPES (pH 7.4), and centrifuged at 15,000 xg for 30 min. The final pellet comprised of synaptosomes was diluted in DPBS and stored at -80°C until further use.

Human synaptosomes were purified from i^3^ LMN after 28 days in culture as described before ^8, 85^. Briefly, i^3^ LMNs were washed twice with DPBS. Then, 2 mL Syn-Per reagent (ThermoFisher Scientific, 87793) was added to each 10-cm cell culture dish. Neurons were scraped, collected, and centrifuged at 1,200 xg for 10 min at 4°C. The supernatant was collected and centrifuged at 15,000 xg for 20 min 4°C. The resulting pellet comprised of synaptosomes was resuspended in DPBS and stored at -80°C until further use.

Mouse and human synaptosomes were labeled with the pH-sensitive dye, pHrodo Red, succinimidyl ester (ThermoFisher Scientific, P36600), or the pH-insensitive dye, Alexa Fluor 546 NHS Ester (AF 546; Invitrogen, A20002) in 100 mM sodium bicarbonate solution (pH 8.5; Sigma, S6297-250G) following the manufacturer’s instructions. Synaptosomes were stored in 5% dimethyl sulfoxide (DMSO; Sigma, D2650) in DPBS at -80°C until further use.

### Live-cell phagocytosis assay and drug treatments

The live-cell phagocytosis assay was performed as described previously in detail^8^. Briefly, PMPs were plated in Primaria 96-well plates (Corning, 353872) and allowed to differentiate into iMGs for 10 days as described^8^. Differentiated iMGs were incubated at 10°C for 10 min and then sonicated pHrodo red-labeled mouse or human synaptosomes were added to the cells. For wells containing iMGs treated with compound, iMGs were pre-treated with 10µM cytochalasin D (Sigma, C8273) or 100nM bafilomycin A1 (Med Chem Express, HY-100558) for 30 min prior to administration of synaptosomes, at which point the drugs were diluted to 5uM and 50nM, respectively. For experiments with rapamycin, iMGs were incubated with 100 nM rapamycin (Combi-Blocks QA-9258) diluted in DMSO for 24h before the assay was conducted; rapamycin was diluted to 50nM upon addition of synaptosomes. iMGs were also treated with 0.05% DMSO as vehicle controls. The plate was centrifuged at 270 xg for 3 min at 10°C to facilitate contact between synaptosomes and iMGs^86^. At least 2h before the assay was conducted, cell nuclei were stained with NucBlue Live Ready reagent (Invitrogen, R37605) as per the manufacturer’s instructions. For phagocytosis of aggregated protein, PFN1 C71G recombinant protein that was described before ^41^ was induced to aggregate by shaking at 1,000 rpm overnight at ambient temperature. The formation of amorphous aggregates was confirmed with transmitted light microscopy using the Leica DMI 6000B inverted fluorescent microscope described above. The aggregated protein was labeled with pHrodo Red and administered to iMGs as described for synaptosomes.

Sixteen phase, blue and red fluorescence images were acquired per well at 20X every hour for 12h using the Cytation 5 cell imaging reader (Biotek). Imaging analysis was performed with Gen5 software (Biotek). For each time point, the area of pHrodo signal was calculated and normalized to the cell number obtained by counting the nuclei in the blue channel. The normalized area for three technical replicates was averaged. For each independent differentiation, the phagocytosis index was calculated with the following equation: normalized area at each timepoint/ (normalized area at 12h for WT iMGs – normalized area at 0h for WT iMGs). The area under the curve was determined using GraphPad Prism v9.3.1. Data from at least three independent differentiations are reported.

### Washout engulfment assay and analysis

iMGs were incubated at 10 °C for 10 min and fed with sonicated AF546-(“0h time point”) or pHrodo red-labeled (“48h post-washout time point”) human synaptosomes. Samples were centrifuged at 270 xg for 3 min at 10°C and then incubated for 15 min at 37°C to allow for initial binding and uptake. Unbound synaptosomes were washed away with DPBS and cells were immediately fixed (0h time point) or incubated at 37°C for an additional 48h with fresh iMG media **Table 1** to allow time for degradation of engulfed material. Immunostaining and fluorescence imaging acquisition was performed as described above.

At the 0h time point, the relative engulfment index was obtained by analyzing individual cells from four independent differentiations. Z-stacks were separated by channel and the background was subtracted from the synaptosome channel by using the “Background subtraction” function with a rolling ball radius of 10 pixels in ImageJ. Since AF546-labeled synaptosomes were used for this time point, it was necessary to exclude unbound synaptosomes that were not washed away from the analysis. To this end, three-dimensional (3D) reconstructions of an overlay of Z-stack images for the IBA1 and synaptosome channels was performed using the 3D Viewer plugin (Fiji, ImageJ). A 3D visualization of the cells allowed for identification of synaptosomes that are located inside the cell volume or in direct contact with the cell boundary defined by IBA1 staining. Synaptosomes that did not meet these criteria were manually deleted from the original Z-stack by an experimentalist blinded to the genotype of the cells. After free synaptosomes were removed, Z-stacks were maximum projected and automatically thresholded using the “Otsu” method for the synaptosome channel and the “Mean” method for the IBA1 channel. The total area of the synaptosomes, as well as the total cell area, were measured by using the “Measure” function in ImageJ. The area of synaptosomes per cell was then normalized to the cell area values for at least 25 cells per condition. Data from the mutant line was normalized to the WT line according to each differentiation.

For the 48h post-washout time point, the volume of pHrodo-labeled synaptosomes located within CD68-positive endolysosomal compartments was measured from individual cells across three independent differentiations. The background signal from the synaptosome (red) channel was subtracted using the “Background subtraction” function described above. Background subtraction was also performed for the CD68 (green) channel, except using a rolling ball radius of 50 pixels. To confirm that the synaptosome signal co-localized with CD68 signal, Z-stacks were maximum projected and both channels were superimposed to visually confirm the presence of yellow (i.e., green overlaid with red) structures. Most of the synaptosome fluorescent signal show spatial overlap with CD68. In rare instances when the signal did not colocalize, the synaptosomes were manually removed from the Z-stacks by an experimentalist blinded to the genotype of the cells. Processed Z-stacks were automatically thresholded using the “Otsu” method and the volume of synaptosomes was measured using the macro “Measure Volume of Thresholded Pixels in an Image Stack” (https://visikol.com/blog/2018/11/29/blog-post-loading-and-measurement-of-volumes-in-3d-confocal-image-stacks-with-imagej/) in ImageJ. The values from at least 10 cells per condition were averaged and normalized to the values of WT iMGs for the corresponding differentiation.

The Cytation 5 cell imaging reader was used to acquire live-cell images of iMGs 48 hours post-synaptosome washout for quantification of the percentage of pHrodo-positive cells remaining at that time point. Two technical replicate wells were imaged per genotype. Twenty FOVs per technical replicate at 20X magnification were imaged. The number of pHrodo-positive cells was manually counted using the Cell Counter plugin (Fiji, ImageJ) by an experimentalist blinded to the genotype of the cells. The total number of cells per FOV was counted similarly using the bright field channel. The sum of the number of cells in all the analyzed FOV was divided to the total number of cells per duplicate and multiplied by 100 to obtain the percentage. The average percentage for technical duplicates was reported. A total of 400 and 600 cells were analyzed per technical replicate across four independent differentiations.

### Stereotactic injection of dead neurons into mice

Primary neurons were prepared from wild-type *C57BL/6* mouse embryos harvested at day E18.5 following our published methods ^87^. Neurons were cultured for nine days before cell death was induced as described before ^35^ with the following modifications. Briefly, neurons were gently washed with DPBS and detached from the plate by gently pipetting DPBS at the bottom of the well. Cell death was induced by exposing neurons to UV light (302nm) for 15 min using a Molecular Imager Gel Documentation System XR+ (Bio-Rad). Dead neurons were maintained on ice for 2-3 hours until the time of stereotactic injection into mice. Neurons were resuspended in 1mL of DPBS followed by the addition of equimolar concentrations of the pH-insensitive dye, Alexa Fluor 546 NHS Ester (AF546), and the pH-sensitive dye, CypHer5E NHS Ester (Cy5E; Cytiva, PA15401) at 37°C for 30 min. Labeled neurons were then diluted in DPBS, harvested by centrifugation at 200 x g for 8 min at 4°C, and washed with DPBS to remove residual dye as per the manufacturer’s instructions. The number of dead cells was determined using Trypan Blue 0.4% staining solution (AMRESCO Inc, K940).

Dead neurons were resuspended in DPBS at a density of 50,000 neurons per microliter for subsequent stereotactic injection into three-month-old Pfn1^em4Lutzy^/J heterozygous mice (referred to as PFN1 C71G^+/-^) and wild-type (WT) littermates. Four PFN1 C71G^+/-^ mice (2 males, 2 females) and five WT control mice (2 males, 3 females) were included in this study. Mice were initially anesthetized through 3% isoflurane inhalation and anesthesia was maintained with 1.5% isoflurane throughout the procedure. Stereotactic injections were performed according to a previously described protocol with the following modifications^88^. Briefly, 1uL containing 50,000 dead neurons was injected into the right hemisphere in the motor cortex at the location of 2 mm in front of bregma and 1 mm right from the midline at the depth of 0.5 mm below the brain surface. As a control, 1uL of saline solution (Bioworld, 40120975-2) was injected contralaterally (2 mm in front of bregma, 1 mm left from the midline, and a 0.5mm depth below the brain surface). Injections were performed at a speed of 200 nL/min using a stereotaxic instrument (KOPF, Model 900LS). Anesthesia was discontinued after the operation was completed. Buprenorphine (1 mg/kg, subcutaneously), cefazolin (500 mg/kg, intramuscularly), meloxicam (5 mg/kg, subcutaneously) were administered prior to the end of surgery session. Animals were placed on top of a heating blanket and their recovery was monitored for ∼15 min and until the mice were ambulatory. After recovery, animals were returned to their home cages for 72h, at which time animals were euthanized with an overdose isoflurane and transcardially perfused with DPBS and 4% paraformaldehyde for subsequent tissue processing.

### Immunohistochemistry, imaging acquisition, and analysis

Mouse brains from the “stereotactic injection of dead neurons” study were dissected and post-fixed with 4% paraformaldehyde overnight, placed in 30% sucrose in 0.1M phosphate buffer (PB) and allowed to sink to the bottom of the conical tube before sectioning. Forty µm coronal sections were prepared, encompassing the entire site of injection as determined by the presence of AF4546 fluorescent signal. The following tissue processing steps were conducted with agitation in a plate rocker at ambient temperature. Floating sections were blocked in 10% goat serum, 0.01% Triton X-100 in 0.1 M PB for 1h. Sections were incubated overnight with rabbit anti-IBA1 antibody at 1:500 (Wako Chemical USA, NC9288364) diluted in blocking buffer at 4°C. Sections were washed and incubated with Alexa fluor-488 secondary antibody (Jackson ImmunoResearch, 711-545-152) at 1:1000 in blocking buffer for 1h and counterstained with DAPI prior to mounting with ProLong Gold anti-fade reagent (Invitrogen, P36930) for subsequent immunofluorescence analysis.

Every tissue section that contained AF4546 fluorescent signal from the site of dead neuron injection was imaged and analyzed by an experimentalist who was blinded to the genotype of the tissues. Z-stack images with a step size of 0.2 µm were acquired with a 20X air objective on a Leica DMI 6000B inverted fluorescent microscope equipped with a Leica DFC365 FX camera and AF6000 Leica Software v3.1.0. Z-stacks were sum projected and the background was subtracted using ImageJ. Images were automatically thresholded using the “Otsu” method in the AF546 channel (constitutively fluorescent dead neurons) and the Cy5E channel (dead neurons in acidic compartments) independently. The area of the signal in each channel was assessed with the “Measure” function in ImageJ. The area of AF546 signal within all AF546-positive brain slices for each mouse was summed, resulting in a total “AF-neuron” area. The same procedure was carried out for Cy5E signal. The total area of the fluorescent signal in Cy5E channel was divided by the total AF-neuron area to determine the fraction of dead neurons located within acidic compartments per mouse.

To quantify the IBA1 signal intensity, an ROI containing scar tissue from the injection site was manually drawn and three concentric rings were automatically generated using an Image J macro (https://www.dropbox.com/s/yp2owicooalm3r1/batch_measureConcentricShapes.ijm?dl=0).The injection site was excluded from analysis. The total signal intensity of each ring was calculated using the “Measure” function (ImageJ) and normalized to the number of IBA1-positive cells counted manually. The values of all the analyzed brain slices per mouse were average and presented as individual data points. All the analyses were performed by a subject blinded to the genotype.

### De-quenched-BSA (DQ-BSA) assay

PMPs were plated in Primaria 24-well plates (Corning, 353847) at a density of 100,000 cells per well for microglia differentiation. iMGs were pulse-labeled with 12 µg/mL green DQ-BSA reagent (Invitrogen, D12050) for 1h, washed with DPBS, and subjected to live-cell imaging after adding fresh iMG media (**Table 1**). As a negative control, cells were treated with 200nM of bafilomycin A for 30 min before the assay. Twenty phase and green fluorescence images were acquired per well from two technical replicates at 20X using the Cytation 5 cell imaging reader (Biotek). Images were acquired immediately after washing and every hour for four hours. Images were analyzed to obtain total DQ-BSA intensity values and the number of cells by counting the cell nuclei from all the acquired fields of view with Gen5 software. Total intensity values were divided normalized to the number of cells and average value of the technical replicates are reported.

### LysoSensor DND-189 staining and analysis

PMPs were plated in Primaria 24-well plates (Corning, 353847) at a density of 100,000 cells per well for microglia differentiation as described above. On day 10, iMGs were loaded with 1 μM LysoSensor Green DND-189 (Invitrogen, L7535), and incubated for 20 min at 37°C. This pH sensor dye produces higher fluorescent signal in acidic environments allowing for a relative comparison of the acidification state in intracellular compartments. Cells were washed twice with PBS and fresh iMG media (**Table 1**) was added before live-cell imaging. At least 2h before the experiment, cell nuclei were stained with NucBlue Live Ready (Invitrogen, R37605) reagent for quantification of the number of cells in each well as per the instructions from the manufacturer. Twenty phase, green and blue fluorescence images were acquired per well from two technical replicates at 20X using the Cytation 5 cell imaging reader (Biotek). Imaging analysis was performed with Gen5 software, where the total green, fluorescent signal intensity was normalized to the cell number. The average of both technical replicates was normalized to the values of WT iMGs for each independent differentiation to obtain the DND-189 relative intensity. Three independent differentiations were used per sample condition. To confirm that the DND-189 relative intensity is dependent on the pH of acidic compartments, lysosomal acidification was inhibited by pre-treating the cells with 200nM of bafilomycin A for 30 min.

### HMC3 culture and PFN1 knockdown

The Human Microglial Clone 3 (HMC3) cell line (CRL-3304) was obtained from ATCC (American Type Culture Collection). HMC3s were maintained in Eagle’s Minimum Essential Media (Corning, 10-009-CV) supplemented with 10% (vol/vol) FBS (Sigma-Aldrich, F4135) under standard culture conditions (37 °C, 5% CO_2_/95% air). Media was changed every 2-3 days. Cells were split with 0.25% trypsin (Gibco, 15090046) when they reached 80-90% confluence and sub-cultured for further passages. For PFN1 knockdown, third-generation lentiviruses carrying a CSCGW2 lentivector plasmid containing miRNA targeting *PFN1* (GCAATAAGGGGTATGGGGTA) or a scramble sequence (TAATCGTATTTGTCAATCAT) under the U6 promoter were produced by VectorBuilder. Cells were transduced at a multiplicity of infection (MOI) of 5 in the presence of 5 µg/mL polybrene. After incubation for 18h, lentiviral particles were removed by changing the media. Cells were maintained for another five days before lysis and western blotting.

### BODIPY staining and quantification of lipid droplets

PMPs were seeded on coverslips and differentiated into iMGs as described above. After 10-12 days of differentiation, half of the media volume was removed and replaced with a staining solution containing BODIPY 493/503 (Invitrogen, D3922) in DMSO, which was further diluted in iMG media (**Table 1**) for a final concentration of 2µM BODIPY 493/503 ^89^. After 15 minutes at 37°C, cells were washed twice with DPBS and fixed with 4% paraformaldehyde for 15 min at ambient temperature. Immunofluorescence staining, imaging acquisition, and deconvolution were performed as mentioned in the section Immunohistochemistry, imaging acquisition, and analysis.

Individual cells from maximum projected images were automatically thresholded by using the “Default” method in ImageJ and the total area of lipid droplets (BODIPY fluorescent signal) was measured using the Particle Analyzer plugin (Fiji, Image). The total area from all cells within a condition (i.e., genotype) were averaged per independent differentiation. For each independent differentiation, data from mutant iMGs were normalized to the corresponding WT iMG condition. A total of 60 cells were analyzed per condition across three independent differentiations.

### Differential scanning fluorimetry (DSF) assay with PIPs

Recombinant PFN1 proteins for DSF were described in a previous study ^41^. The 1,2-dioctanoyl-sn-glycero-3-phospho-(1’-myo-inositol-4’,5’-bisphosphate) 08:0 (PIP2; Avanti Polar Lipids, 850185P) and phosphatidylinositol 3-phosphate diC8 (PI3P; Echelon Biosciences, P-3008) were dissolved in PBS to the different concentrations (20µM, 50µM and 200µM)and mixed with 25X SYPRO Orange (Invitrogen #S6651). Purified WT or mutant PFN1 proteins were added to each sample tube to a final concentration of 20µM and plated in duplicate in 384-well plates (Bio-Rad, HSR-4805). The samples were subjected to heat denaturation using CFX384 Touch Real-Time PCR Detection System. The temperature gradually increased from 25°C to 70°C, with 0.3°C increments every 5 sec. The fluorescence intensity was acquired at each time point with a HEX detector (excitation 515–535 nm, emission 560–580 nm). The fluorescence intensity values from duplicate samples were averaged and plotted as a function of temperature to produce melting curves in GraphPad Prism. Melting curves were fit with a Boltzmann’s sigmoidal function in GraphPad Prism to determine the apparent melting temperature (T_m_). The difference between the T_m_ of each experimental group and the no-lipid control condition was reported as the ΔT_m_. At least three independent experiments were performed per condition.

### Recombinant PFN1 Protein Expression for NMR experiments

WT and PFN1 M114T variants were cloned into a modified pet28 vector with a SUMO tag between BamHI restriction site. Both WT and M114T variants were expressed in BL21(DE3) *Escherichia coli* competent cells and isotopically labelled with ^15^N by growing the cells in M9 media containing 1 g of ^15^NH_4_Cl per L. The cells were grown at 37 °C to an OD_600_ of 0.8 and then induced using 1 mM isopropyl-β-d-1-thiogalactopyranoside (IPTG; GoldBio, 2481C) for 8 h at 30 °C. Cells were harvested and lysed by sonication in 40 mL of buffer containing 50 mM Tris HCl pH 8, 300 mM NaCl, 1 mM Tris (2-carboxyethyl) phosphine (TCEP; GoldBio, TCEP25), and 25 mM imidazole (Fischer Scientific, BP-305-50). Lysates were centrifuged at 20,000 rpm at 4 °C for 1 h and passed through a pre-equilibrated 15 mL prepacked Histrap^TM^ HP nickel column (Cytiva, 17524802), washed with 5 column volumes of 50 mM Tris HCl pH 8, 300 mM NaCl, 1 mM TCEP and 25 mM imidazole, and eluted with 50 mM Tris HCl pH 8, 300 mM NaCl, 1 mM TCEP, and 300 mM imidazole. The SUMO tag was cleaved off with 5 mL Ubiquitin-like-specific protease 1 (ULP1) at a 1:5 ULP1 to protein ratio during an overnight dialysis at 4 °C^90^. The protein is then passed through a second round of nickel column purification. The purified protein was buffer exchanged into 50 mM phosphate buffer pH 6.5, 150 mM NaCl, 200 μM TCEP, 1 mM EDTA, and 50 mM L-Arg and L-Glu by overnight dialysis then concentrated using a 10 kDa Centriprep concentrator (Millipore, UFC901024). The concentrated protein was passed through a pre-equilibrated size exclusion column (Cytiva Custom Superdex ^TM^ 75 Increase HiScale ^TM^ 16/40, CP 19-186)

### NMR Binding Studies

Binding of PFN1 WT and M114T to PI3P (described above in the DSF experiments) was monitored via NMR spectroscopy. ^15^N-^1^H heteronuclear single-quantum coherence (HSQC) spectra were collected at 293 K on a Varian Inova spectrometer (Palo Alto, CA) operating at 600 MHz equipped with a triple-resonance cold probe. Data processing was performed using NMRPipe^91^ and Sparky software^92^. Uniformly ^15^N labelled PFN1 WT and M114T mutant were dissolved in 600 μL of buffer (2 mM HEPES, 0.2 mM CaCl_2_, 0.2mM ATP, 50 mM L-Arg, and 50 mM L-Glu at pH 7, 92% H_2_O/8% ^2^H_2_O) at initial concentrations of 50μM. PI3P was dissolved in the same buffer to achieve a stock concentration of 2.7 mM. A titration was performed at 20° C by adding increasing amounts of PI3P to reach the following concentrations: 10 μM, 20 μM, 30 μM, 40 μM, 50 μM, 67.5 μM, 85 μM, 100 μM, 125 μM, 150μM, 175μM, 202 μM, 250 μM, 350 μM, 500 μM, 750 μM, 1100 μM, 1450 μM for PFN1 WT and 10 μM, 20 μM, 30 μM, 40 μM, 50 μM, 67.5 μM, 85 μM, 100 μM, 125 μM, 150μM, 200 μM, 250 μM, 350 μM, 500 μM, 750 μM, 1100 μM, 1450 μM for PFN1 M114T. A ^15^N-^1^H HSQC spectrum was collected after each addition of PI3P. The chemical shift perturbations were calculated for each PI3P titration point according to the following equation:

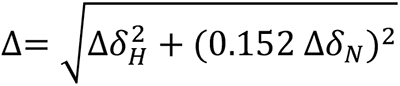

where Δδ*_H_* and Δδ*_N_* are the ^1^H_N_ and ^15^N chemical shift differences measured for each residue in each ^15^N-^1^H HSQC spectrum.

The apparent *K*_d_ was derived from global fitting of the following residues: Ala 5, Tyr 6, Ile 7, Asn 9, Asp 13, Gln 17, Asp 18, Trp 31, Lys 104, Lys 107, Tyr 128, Glu 129, Met 130, Ala 131, Leu 134 for PFN1 WT, and Ala 5, Tyr 6, Ile 7, Asp 13, Gln 17, Asp 18, Ser 56, Ser 57, Glu 129, Met 130 and Leu 134 for PFN1 M114T. The residue specific titration curves were fitted using the following equation to determine K_d_ using Matlab^93^,

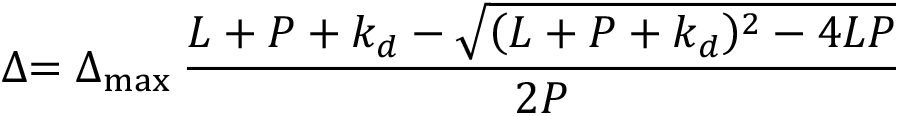

where Δ_max_ is the maximum chemical shift perturbation, L and P are the total ligand and protein concentrations and K_d_ is the apparent dissociation constant.

### Statistical analysis

Statistical analyses were calculated using GraphPad Prism v9.3.1. Statistical tests are stated in the figure legends. The number of independent experiments is stated in the corresponding methods section and in the figure legends.

## Acknowledgements

We are grateful to Dr. Michael Ward (NINDS, MD) for providing the WTC11 iPSC line with the hNIL cassette, Dr. Jeffery Kelly (the Scripps Research Institute, CA) and his lab for providing Rapamycin and advice, Drs. Yen-Chen Lin (UMass Chan) and Desiree Baron (UMass Chan) for help with iPSC culture, Dr. Anthony Giampetruzzi (UMass Chan) for help with the Cytation 5, Dr. Travis Faust for advice on mouse studies, the UMass Chan Mass spectrometry Facility for the proteomics experiments, and the UMass Chan Transgenic Animal Modeling Core for in vitro fertilization of mutant PFN1 mice, the Cellular Engineering Service at The Jackson Laboratory for expert assistance with gene-editing of iPSCs described in this manuscript. D.A.B. is supported by the NIH/NINDS (R01 NS108769, R21 NS120126), NIH/NIGMS (R01 GM137529, R01 GM147677), the Department of Defense (W81XWH202071/PRARP), the Angel Fund for ALS research, the Radala Foundation and the Robert Packard Center for ALS Research. This project has also been supported by the Dan and Diane Riccio Fund for Neuroscience (UMass Chan; to D.A.B. and D.P.S.) and NIH/NIGMS R01GM137529 (F.M. and D.A.B.). We are also grateful for the following support: NIMH-R01MH113743 (D.P.S.); NINDS-R01NS117533 (D.P.S.); NIA-RF1AG068281(D.P.S.); U54OD020351 (to the Jackson Laboratory Center for Precision Genetics, C.L.)

