## Supplemental Figures for "Expression of ALS-PFN1 impairs vesicular degradation in iPSC-derived microglia"

### SUPLEMMENTARY FIGURES

a

| | Code | Line Name | Genotype |
|---|---|---|---|
| C71G-PFN1 isogenic group | WT | KOLF2.1J: PFN1_C71G WT clone B03 | WT clone during C71G mutation editing |
| | C71G$^{+/-}$ | KOLF2.1J: PFN1_C71G het clone C01 | C71G/WT |
| M114T-PFN1 isogenic group | WT | KOLF2.1J: PFN1_M114T WT clone H09 | WT clone during M114T mutation editing |
| | M114T$^{+/-}$ | KOLF2.1J: PFN1_M114T het clone A08 | M114T/WT |
| | M114T$^{+/+}$ | KOLF2.1J: PFN1_M114T homo clone D12 | M114T/M114T |

b

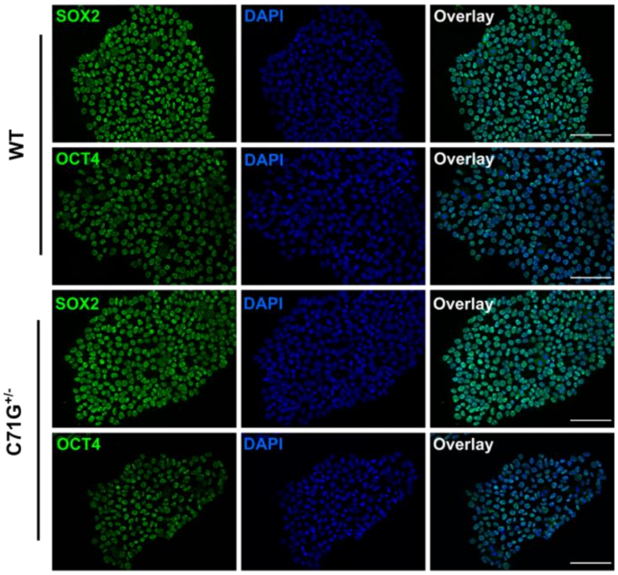


c

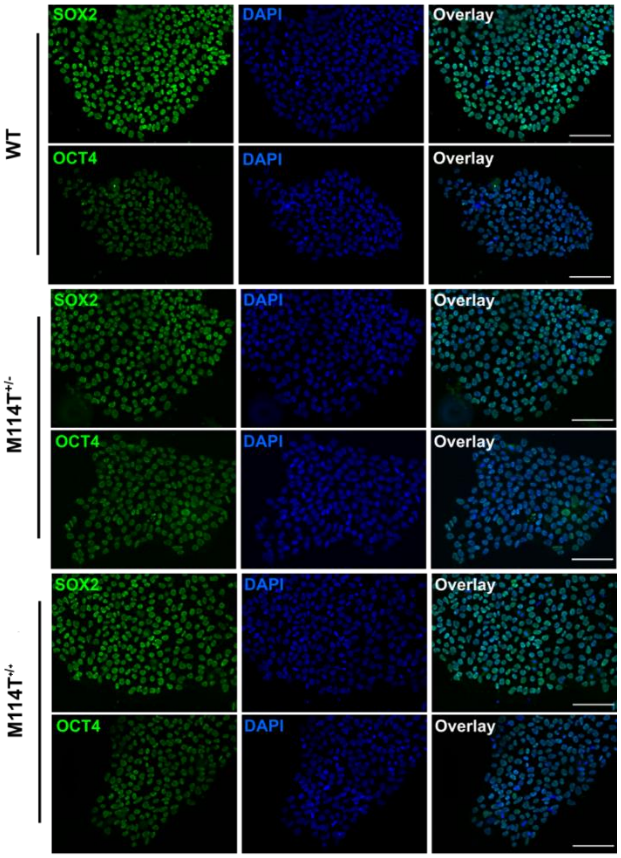


**Supplemental Figure 1. Pluripotency analysis of iPSC lines used in this study. a,** The iPSC lines used in this report. **b-c,** Immunofluorescence images of pluripotency markers SRY-Box transcription factor 2 (SOX2; green) and octamer-binding transcription factor 4 (OCT4; green) in WT and C71G$^{+/-}$ iPSCs (**b**) and WT, M114T$^{+/-}$ and M114T$^{+/+}$ iPSCs (**c**). Representative images obtained from one analysis. Scale bar: 100 µm.

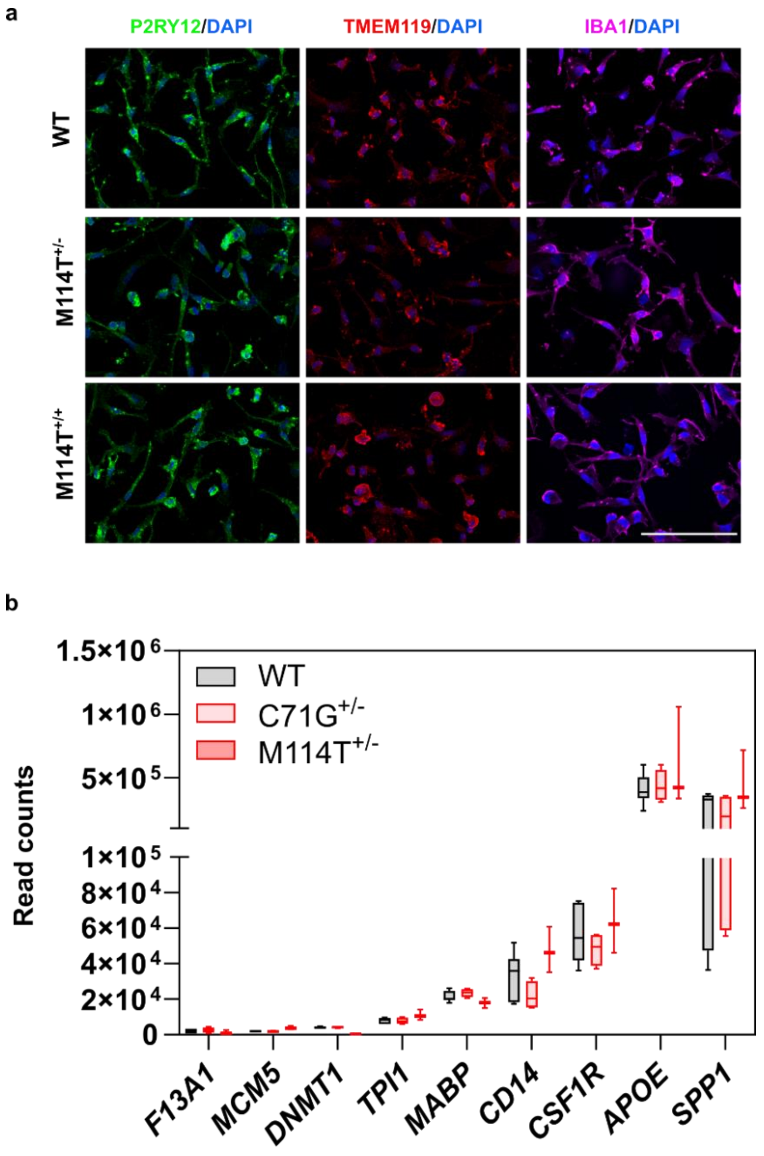


**Supplemental Figure 2. Expression of microglia and myeloid markers in ALS-PFN1 and isogenic control iMGs. a**, Representative immunofluorescence images of the microglia and myeloid markers P2RY12, transmembrane protein 119 (TMEM119), and ionized calcium-binding adapter molecule 1 (IBA1) in WT, M114T$^{+/-}$ and M114T$^{+/+}$ iMGs. Scale bar: 100 µm. **b,** Normalized read counts of genes associated with different microglia developmental stages expressed in mutant PFN1 and WT iMGs, including early-stage yolk-sac microglia progenitor gene *F13A1* and embryonic-microglia genes minichromosome maintenance complex component 5 (*MCM5*), DNA methyltransferase 1 (*DNMT1*), and triosephosphate isomerase 1 (*TPI1*) are shown. Also shown are adult-microglia genes V-Maf musculoaponeurotic fibrosarcoma oncogene homolog B (*MAFB*), monocyte differentiation antigen CD14 (*CD14*) and colony-stimulating factor-1 receptor *(CSFR1)*, and the aging-related microglia genes Apolipoprotein E (*APOE)* and secreted phospho-protein 1 (*SPP1).* Box and whisker plots of WT n=7, C71G$^{+/-}$ n=4, and M114T$^{+/-}$ n=3 independent differentiations.

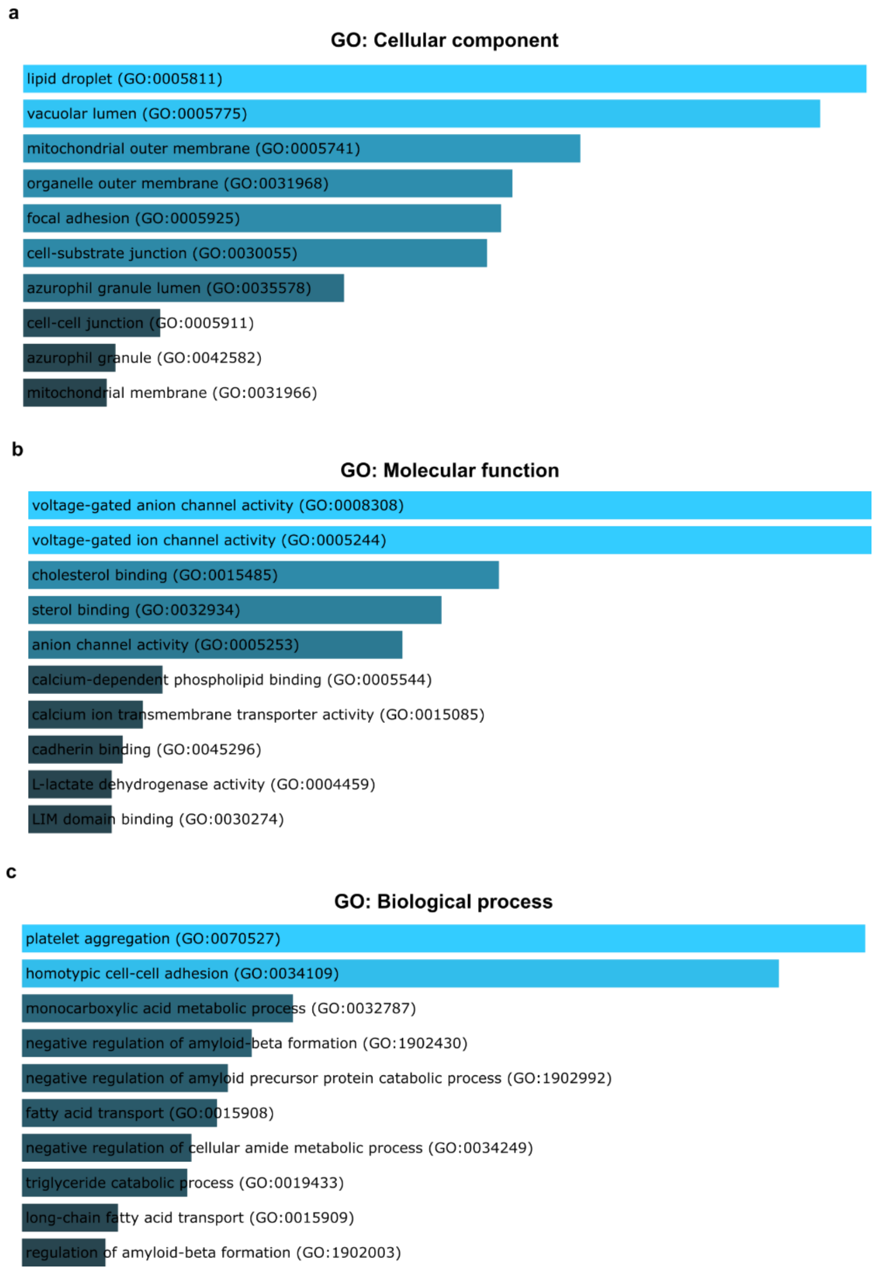


**Supplemental Figure 3. Gene ontology (GO) analysis of differentially expressed proteins from the proteomics analysis of C71G$^{+/-}$ versus WT iMGs. a-c,** Heat maps obtained by Enrichr showing the top 10 GO terms for three different GO groups: cellular component (**a**), molecular function (**b**), and biological process (**c**). The bars are colored according to *P* value. Additional terms and statistical values can be found in **Table S4.**

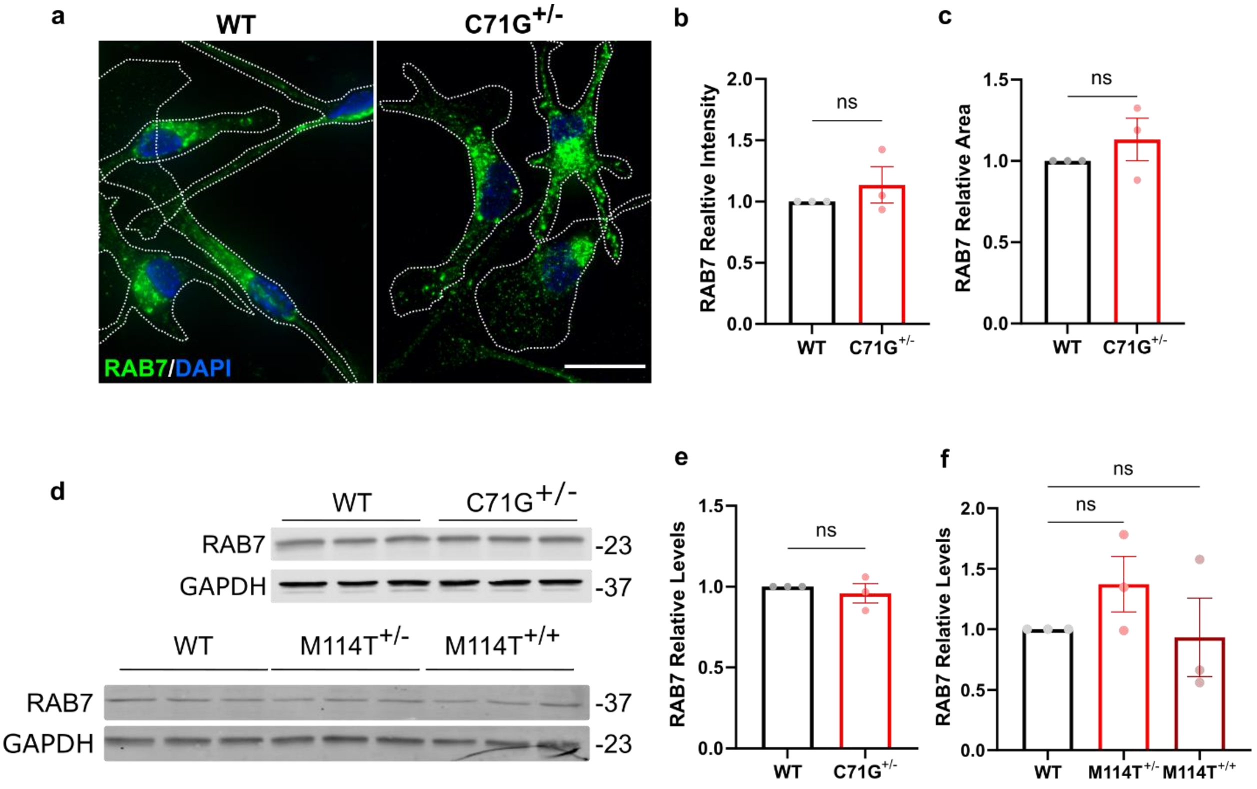


**Supplemental Figure 4. RAB7 expression is similar among WT and ALS-PFN1 iMGs. a-c** Immunofluorescence analysis of RAB7 in WT and PFN1 C71G$^{+/-}$ iMGs (n=3 independent differentiations). **a,** Representative immunofluorescent images. Cell boundaries defined by anti-PFN1 staining (not shown for clarity) are depicted with white dashed lines. Scale bar=50 µm. **b,c,** Quantification of the mean intensity (**b,** ns *P*=0.4094, t=0.9205, df=4) and area (**c,** ns *P*= 0.3701, t=1.009, df=4) of RAB7 immunofluorescence signal analyzed on a per cell basis. Mutant iMG data were normalized to the corresponding WT condition within the same experimental differentiation. **d,** Western blot analysis of RAB7 with GAPDH used as a loading control for cell lysates derived from C71G$^{+/-}$, M114T$^{+/-}$, M114T$^{+/+}$ iMGs and their respective WT iMG counterpart (n=3 independent differentiations). **e**, Quantification of **d** for WT and C71G$^{+/-}$ iMGs (**e,** ns *P*=0.5405, t=0.6683, df=4). **f,** Quantification of **d** for WT, M114T$^{+/-}$ and M114T$^{+/+}$ iMGs (**f,** ns *P*=0.2518, q= 1.633, DF=6 for WT vs M114T$^{+/-}$ and *P*= 0.7486, q= 0.6546, DF=6 for WT vs M114T$^{+/+}$). **e,f,** For each independent differentiation, RAB7 protein levels were normalized to the levels of the respective WT controls from the same experimental differentiation. All graphs show mean ± SEM with individual data points representing independent differentiations. Statistics were determined using unpaired two-tailed t-test for WT vs C71G$^{+/-}$ iMGs comparisons and ordinary one-way ANOVA with Dunnett's multiple comparisons test for WT vs M114T$^{+/-}$ and M114T$^{+/+}$ comparisons.

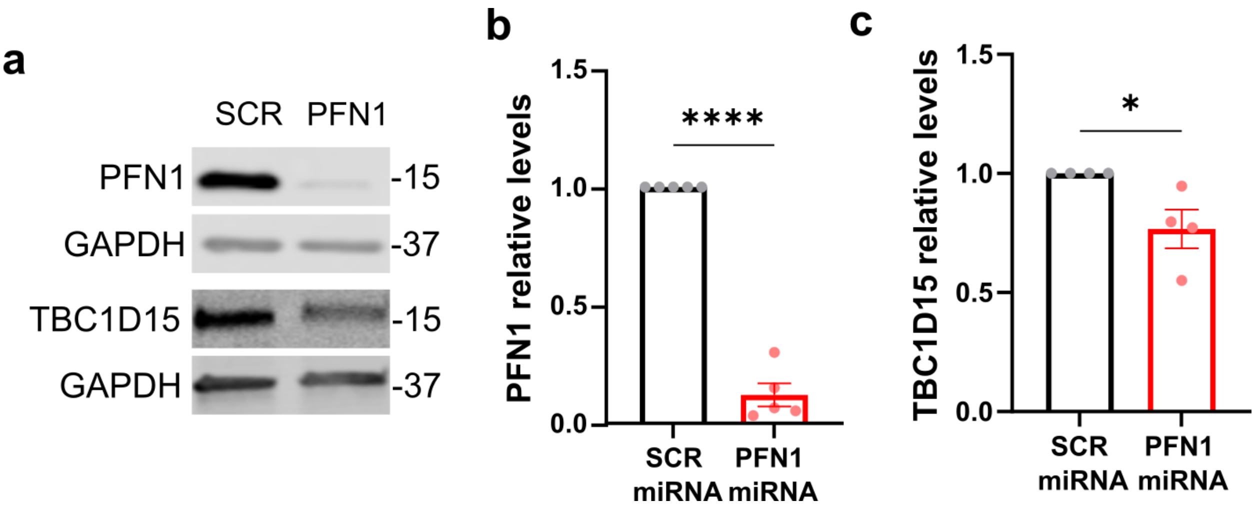


**Supplemental Figure 5. PFN1 knockdown in HMC3 microglia cells correlates with reduced TBC1D15 levels.** PFN1 knockdown in HMC3 cells using miRNAs targeting *PFN1* or a scrambled (SCR) sequence for n=5 independent experiments. **a,** Representative Western blot of PFN1, TBC1D15 and GAPDH, used as loading control. **b,** Quantification of PFN1 (****P=<0.0001, t=17.82, df=8) levels from Western blots exemplified in **a**. **c,** Quantification of TBC1D15 (*P*=0.0289, t=2.857, df=6) as described in **b**. **b,c,** The indicated protein was normalized to GAPDH and to SCR controls for each independent experiment. All graphs show mean ± SEM. Data points in bar graphs represent individual experiments. Statistics were performed by unpaired two-tailed t-test.

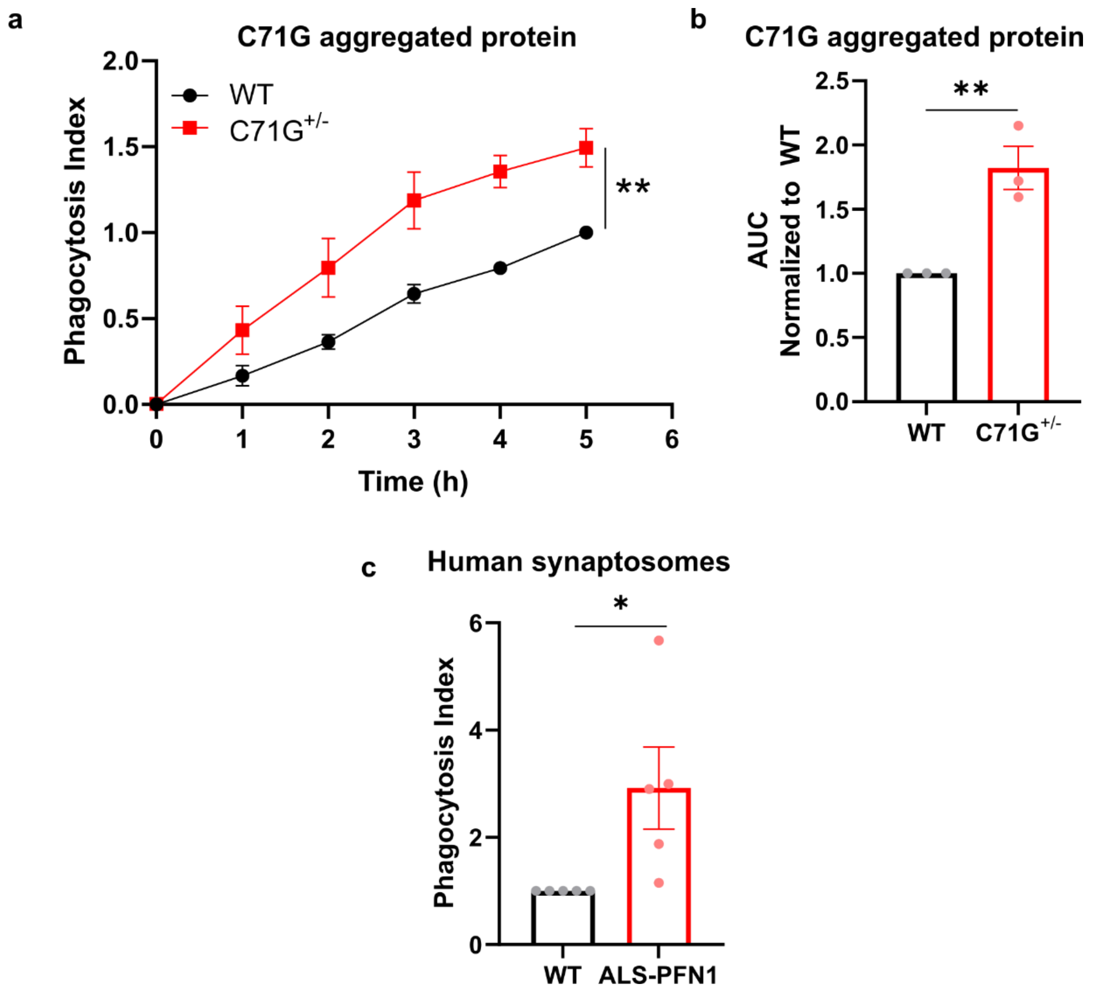


**Supplemental Figure 6. Differential phagocytosis indices for ALS-PFN1 and WT iMGs upon administration of disease-relevant substrates. a-c,** Live-cell phagocytosis assays using pHrodo-labeled disease-relevant substrates. **a,** Quantification of the phagocytosis index (see Methods) for WT and C71G$^{+/-}$ iMGs (n=3 independent differentiations) in the presence of pHrodo-labeled PFN1 C71G aggregated purified protein over a time course of 5h (paired two-tailed t-test, ***P*= 0.0072, t=4.376, df=5). **b,** Area under the curve (AUC) determined from **a** (unpaired two-tailed t-test, ***P*= 0.0082, t=4.881, df=4). **c**, Quantification of the phagocytosis index for C71G$^{+/-}$ (n=3 independent differentiations) and M114T$^{+/-}$ (n=2 independent differentiations), referred to collectively as "ALS-PFN1", and their respective WT controls (n=5 independent differentiations) in the presence of human synaptosomes after 2h of phagocytosis (unpaired two-tailed t-test, **P*= 0.0369, t=2.501, df=8). Mutant iMG data was normalized to the levels of the respective WT control from the same experimental differentiation. All graphs show mean ± SEM with individual data points representing independent differentiations.

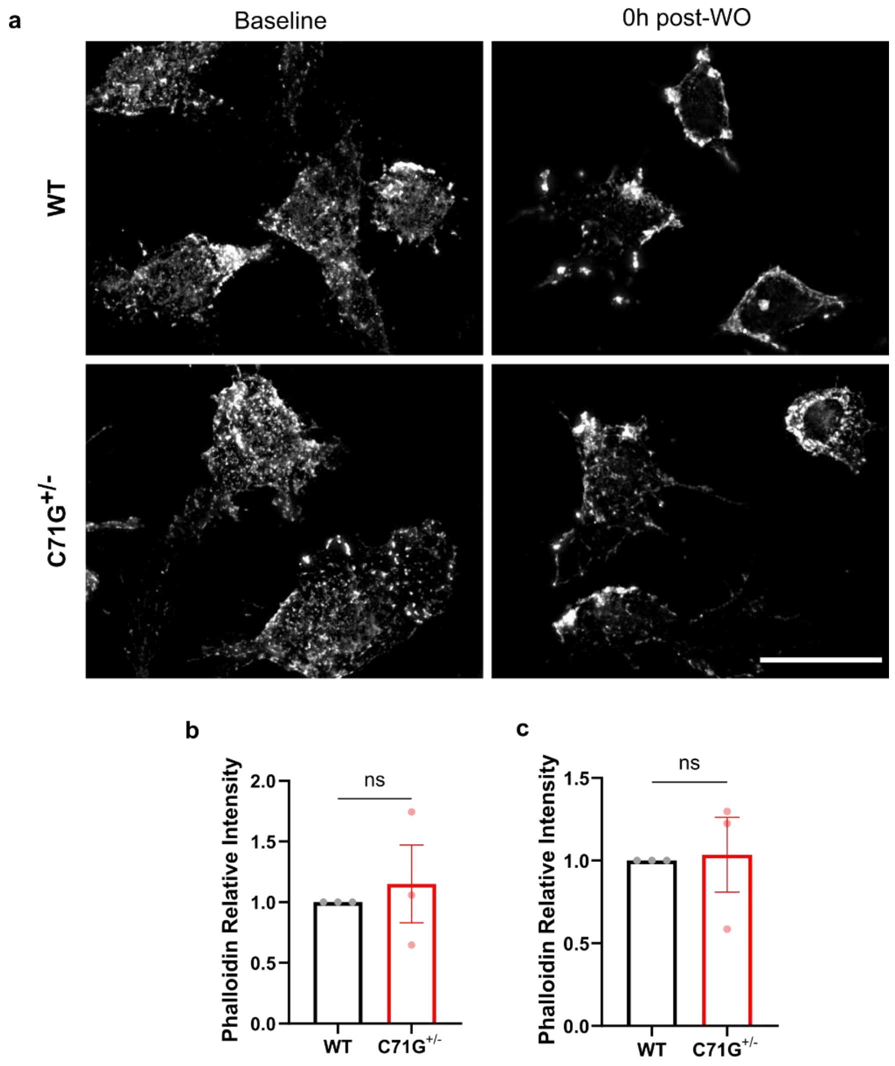


**Supplemental Figure 7. Analysis of F-actin detected by phalloidin staining of PFN1 C71G$^{+/-}$ and WT iMGs subjected to the live-cell phagocytosis assay. a,** Representative immunofluorescence images of phalloidin staining of PFN1 WT and C71G$^{+/-}$ iMGs before administration of human synaptosomes (baseline) and immediately after synaptosomes were washed out (0h post-WO). Scale bar: 25 µm **b,c,** Quantification of phalloidin mean fluorescent intensity. Data from C71G$^{+/-}$ iMGs was normalized to that of WT iMGs within the same experimental differentiation. All graphs show mean ± SEM with individual data points representing n=3 independent differentiations. Unpaired two-tailed t-test was used for all statistical comparisons (**b,** ns *P*=0.8882, t=0.1591, df=2 and **c,** ns *P*=0.6621, t=0.4711, df=4).

a

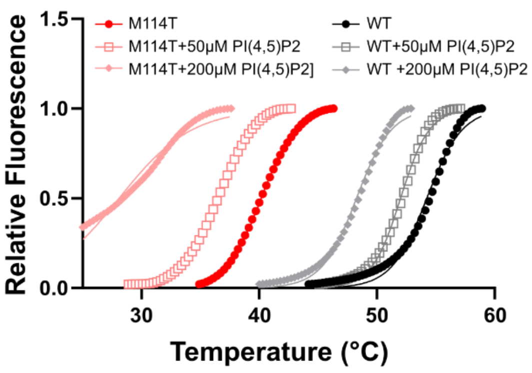


c

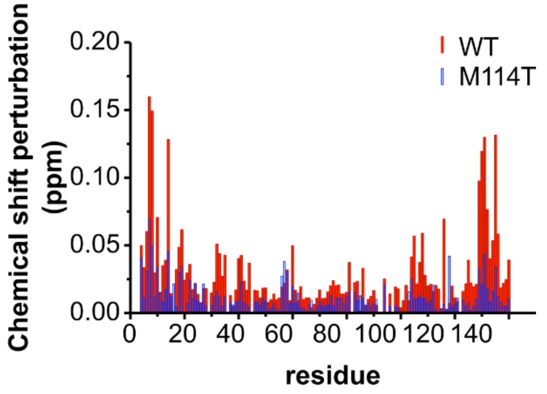


b

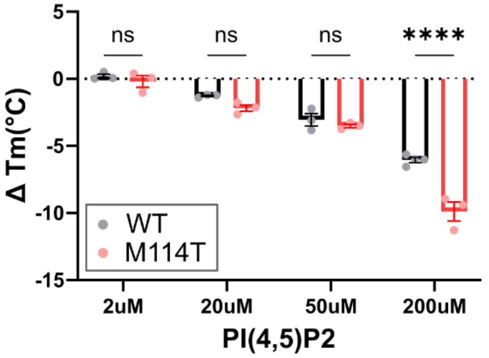


d

| SSE | Residues |
|---|---|
| $\alpha_1$ | 4-12 |
| $\beta_1$ | 17-24 |
| $\beta_2$ | 30-34 |
| $3_{10}$ | 38-41 |
| $\alpha_2$ | 45-52 |
| $\alpha_3$ | 56-62 |
| $\beta_3$ | 64-66 |
| $\beta_4$ | 69-77 |
| $\beta_5$ | 85-90 |
| $\beta_6$ | 99-105 |
| $\beta_7$ | 109-115 |
| $\alpha_4$ | 121-138 |

**Supplemental Figure 8. Additional data pertaining to binding of purified PFN1 variants with phosphoinositides. a,** Thermal denaturation profiles of PFN1 proteins incubated with different concentrations of PI(4,5)2P measured by SYPRO Orange fluorescence as a function of increasing temperature as described for main text **Figure 8**. An average of two technical replicates is shown, which are representative of n=3-4 independent experiments. The curves were fit with the Boltzmann's sigmoidal function to determine an apparent melting temperature ($T_m$). **b**, $\Delta T_m$ reflects the difference between the $T_m$ of the PFN1 variant with the indicated concentration of PI(4,5)2P and the $T_m$ of that PFN1 variant without PI(4,5)2P. Statistics were determined using two-way ANOVA F (1, 28) = 7.115 and Šídák's multiple comparisons test (*****P*=<0.0001 and ns *P*=0.2851, 0.7847). Bar graph shows mean ± SEM with each data point representing an independent experiment. **c,d,** Additional information for the NMR titration studies of PFN1 with PI3P as described in main text **Figure 8** **c,** Chemical shift perturbation (ppm) for each residue of PFN1 between the free and the PI3P-bound state for PFN1 WT (red lines) and PFN1 M114T (blue lines). The secondary structural elements (SSE) formed by the indicated PFN1 sequences are shown and defined as per [25].

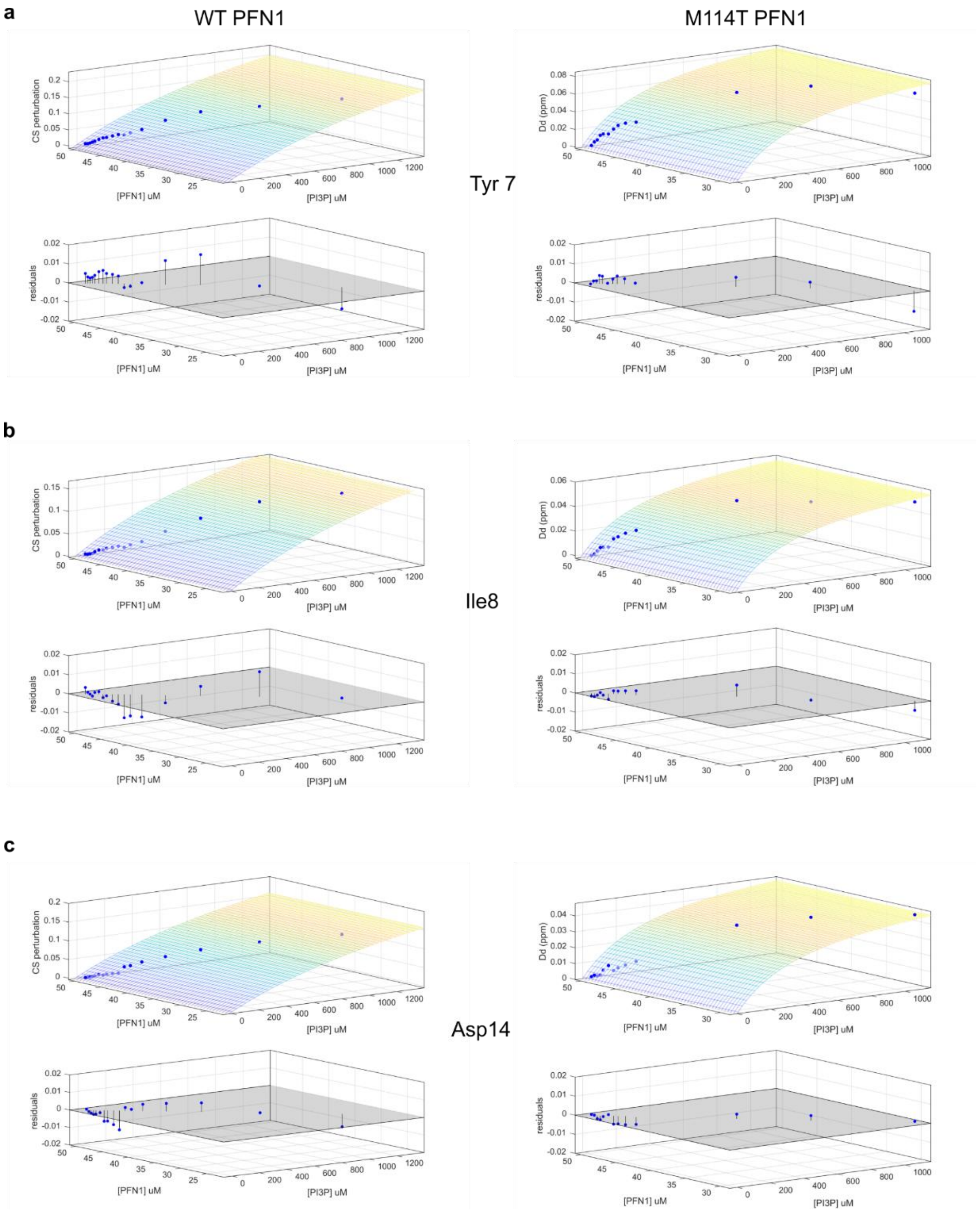


**Supplemental Figure 9. Representative residuals of the MATLAB fits for the NMR titration data for PFN1 with PI3P.** The change in chemical shift (CS perturbation in ppm) for the indicated PFN1 residue (Tyr 7, Ile8 and Asp14) is plotted as a function of PFN1 concentration (uM) and PI3P concentration for both PFN1 WT (left column) and PFN1 M114T (right column). The resultant titration curves were fitted in MATLAB to obtain a dissociation constant as described in the methods. Residuals of the fitting are shown in blue.

### SUPPLEMENTARY TABLES

**Table S1. Additional statistical analysis for this study.** The table contains results for statistical analysis in: **Figure 1c**, statistics for C71G$^{+/-}$ cells (iMGs vs PMPs and iPSCs) using two-way ANOVA and Šídák's multiple comparisons test. **Figure 1d**, additional comparisons not listed in the main text figure for WT iMGs and all comparisons for PFN1 C71G$^{+/-}$ iMGs, including LPS treated versus untreated conditions, using two-way ANOVA and Šídák's multiple comparisons test. **Figure 3b and d**, the indicated comparison was assessed by two-way ANOVA followed by Tukey's multiple comparisons for WT vs C71G$^{+/-}$ iMGs (**b**) and WT vs M114T$^{+/-}$ vs M114T$^{+/-}$ iMGs (**d**).

**Table S2. Results of the differential expression analysis of C71G$^{+/-}$ vs WT iMGs using quantitative proteomics.** Data resulting from the mass spectrometry experiment using tandem mass tag (TMT) quantitative proteomics to identify differentially expressed proteins between PFN1 WT and C71G$^{+/-}$ iMGs. PFN1 WT iMGs was used as the reference condition. Data was analyzed using Scaffold Software as described in the methods. Statistics were determined using T-test and Benjamin-Hochberg correction for multiple testing. Differentially expressed proteins in this table have a *P*- value<0.00160 and are considered statistically significant with the Benjamin-Hochberg test.

**Table S3. Functional enrichment analysis of differentially expressed proteins identified from the quantitative proteomics study.** The top 20 clusters with their representative enriched terms (one per cluster) from pathway and process enrichment analysis using Metascape. Differentially expressed proteins used for this analysis are listed in **Supp. Table 2**.

**Table S4. Gene ontology (GO) analysis of differentially expressed proteins from the quantitative proteomics study.** Enrichr was used to determine enriched GO terms for biological process, molecular function, and cellular component categories from the differentially expressed proteins listed in **Supp. Table 2**. The results of the analysis for each category are presented in different tabs.

**Table S5. Results of differential gene expression analysis from the RNASeq dataset.** RNAseq data obtained from C71G$^{+/-}$ (CG) and M114T$^{+/-}$ (MThet) iMGs (ALS-PFN1 group) and WT controls (WT and WT2) were analyzed using DESeq2 package and Wald test.
