## Supplemental Table 1 for "Expression of ALS-PFN1 impairs vesicular degradation in iPSC-derived microglia"

| **Figure 1c** | | | | |
| --- | --- | --- | --- | --- |
| **Gene** | **Comparison for C71G^+/-^ cells** | ***P value*** | | |
| *PROS1* | iPSCs vs. iMGs | 0.0032 | | |
|  | PMPs vs. iMGs | 0.0212 | | |
| *GPR34* | iPSCs vs. iMGs | 0.0035 | | |
|  | PMPs vs. iMGs | 0.0124 | | |
| *P2RY12* | iPSCs vs. iMGs | 0.0019 | | |
|  | PMPs vs. iMGs | 0.0019 | | |
| *MERTK* | iPSCs vs. iMGs | 0.0013 | | |
|  | PMPs vs. iMGs | 0.0047 | | |
| *SPI-1* | iPSCs vs. PMPs | 0.0209 | | |
| *SOX2* | iPSCs vs. iMGs | <0.0001 | | |
| **Figure 1d** | | | | |
| **Cytokine** | **Other comparison for WT iMGs** | ***P value*** | | |
| IL-6 | Untreated vs 6h LPS | 0.3593 | | |
| IL-10 | Untreated vs 6h LPS | 0.3375 | | |
| CCL5 | Untreated vs 6h LPS | 0.9907 | | |
| TNF-α | Untreated vs 24h LPS | 0.9964 | | |
| **Cytokine** | **Comparison for C71G^+/-^ iMGs** | ***P value*** | | |
| IL-6 | Untreated vs 6h LPS | 0.4469 | | |
|  | Untreated vs 24h LPS | 0.0146 | | |
| IL-10 | Untreated vs 6h LPS | 0.2287 | | |
|  | Untreated vs 24h LPS | 0.0018 | | |
| CCL5 | Untreated vs 6h LPS | 0.9832 | | |
|  | Untreated vs 24h LPS | 0.0076 | | |
| TNF-α | Untreated vs 6h LPS | 0.0125 | | |
|  | Untreated vs 24h LPS | 0.9967 | | |
| **Figure 3b** | | | | |
| ***Comparison*** | | ***P value*** | ***q*** | ***DF*** |
| WT vs. C71G^+/-^ | | <0.0001 | 17.28 | 6 |
| WT vs. WT CytoD | | <0.0001 | 16.69 | 6 |
| C71G^+/-^ vs. C71G^+/-^ CytoD | | <0.0001 | 31.04 | 6 |
| WT CytoD vs. C71G^+/-^ CytoD | | 0.0965 | 1.018 | 6 |
| **Figure 3d** | | | | |
| ***Comparison*** | | ***P value*** | ***q*** | ***DF*** |
| WT vs. M114T^+/-^ | | <0.0001 | 11.29 | 9 |
| WT vs. M114T^+/+^ | | <0.0001 | 10.71 | 9 |
| M114T^+/-^ vs. M114T^+/+^ | | 0.9999 | 0.5756 | 9 |
| WT vs. WT BafA | | <0.0001 | 14.08 | 9 |
| M114T^+/-^ vs. M114T^+/-^ BafA | | <0.0001 | 24.07 | 9 |
| M114T^+/+^ vs. M114T^+/+^ BafA | | <0.0001 | 23.42 | 9 |
| WT BafA vs. M114T^+/-^ BafA | | >0.9999 | 0.09116 | 9 |
| WT BafA vs. M114T^+/+^ BafA | | >0.9999 | 0.2147 | 9 |
| M114T^+/-^ BafA vs. M114T^+/+^ BafA | | >0.9999 | 0.1235 | 9 |
